# Dystroglycan Maintains Inner Limiting Membrane Integrity to Coordinate Retinal Development

**DOI:** 10.1101/121756

**Authors:** Reena Clements, Rolf Turk, Kevin P. Campbell, Kevin M. Wright

## Abstract

Proper neural circuit formation requires the precise regulation of neuronal migration, axon guidance and dendritic arborization. Mutations affecting the function of the transmembrane glycoprotein dystroglycan cause a form of congenital muscular dystrophy that is frequently associated with neurodevelopmental abnormalities. Despite its importance in brain development, the role for dystroglycan in regulating retinal development remains poorly understood. Using a mouse model of dystroglycanopathy (*ISPD^L79*^*) and conditional *dystroglycan* mutants of both sexes, we show that dystroglycan is critical for the proper migration, axon guidance and dendritic stratification of neurons in the inner retina. Using genetic approaches, we show that dystroglycan functions in neuroepithelial cells as an extracellular scaffold to maintain the integrity of the retinal inner limiting membrane (ILM). Surprisingly, despite the profound disruptions in inner retinal circuit formation, spontaneous retinal activity is preserved. These results highlight the importance of dystroglycan in coordinating multiple aspects of retinal development.

**Significance Statement:** The extracellular environment plays a critical role in coordinating neuronal migration and neurite outgrowth during neural circuit development. The transmembrane glycoprotein dystroglycan functions as a receptor for multiple extracellular matrix proteins, and its dysfunction leads to a form of muscular dystrophy frequently associated with neurodevelopmental defects. Our results demonstrate that dystroglycan is required for maintaining the structural integrity of the inner limiting membrane (ILM) in the developing retina. In the absence of functional dystroglycan, ILM degeneration leads to defective migration, axon guidance and mosaic spacing of neurons, and a loss of multiple neuron types during retinal development. These results demonstrate that disorganization of retinal circuit development is a likely contributor to visual dysfunction in patients with dystroglycanopathy.

## Introduction

The precise lamination of neurons is critical for establishing proper connectivity in the developing nervous system. The retina is organized in three cellular layers: the outer nuclear layer (ONL) comprised of rod and cone photoreceptors; the inner nuclear layer (INL) containing horizontal cells, bipolar cells and amacrine cells; and the ganglion cell layer containing RGCs and displaced amacrine cells (Bassett and Wallace, 2012). Two synaptic lamina form postnatally: the outer plexiform layer (OPL), which contains synapses between photoreceptors, bipolar cells and horizontal cells; and the inner plexiform layer (IPL), which contains synapses between bipolar cells, amacrine cells and RGCs. The molecular cues that direct the laminar positioning of neurons and the stratification of their processes within the synaptic layers remain poorly understood.

Unlike the cerebral cortex, where many neurons migrate along the radial glia scaffold, retinal migration does not require contact between neurons and neuroepithelial cells (Reese, 2011). RGCs, bipolar cells, and photoreceptors migrate by nuclear translocation through a basally-directed process. Basal process contact with the inner limiting membrane (ILM) is critical for the polarization and migration of RGCs (Randlett et al., 2011). The ILM is enriched with extracellular matrix (ECM) proteins including laminins, Collagen IV, and perlecan (Taylor et al., 2015; Varshney et al., 2015). Mutations in specific laminins (Lamα1, Lamβ2 and Lamγ3) or the laminin receptor β1-Integrin disrupt formation of the ILM and organization of the ganglion cell layer (GCL) (Edwards et al., 2010; Pinzon-Duarte et al., 2010; Gnanaguru et al., 2013; Riccomagno et al., 2014). How laminins and other ECM proteins are initially organized in the ILM and how the ILM directs the organization of the retina remains unclear.

In addition to β1-Integrin, the transmembrane glycoprotein dystroglycan functions as a receptor for laminins and other ECM proteins through its extracellular α-subunit. Dystroglycan connects to the actin cytoskeleton through the intracellular domain of its transmembrane β-subunit, which is part of the dystrophin glycoprotein complex (Moore and Winder, 2010). Mutations disrupting the glycosylation of dystroglycan affect its binding to Laminin G (LG)-domain containing ECM proteins and lead to a form of congenital muscular dystrophy termed dystroglycanopathy (Taniguchi-Ikeda et al., 2016). The most severe forms, Muscle-Eye-Brain disease (MEB) and Walker Warburg Syndrome (WWS), are accompanied by cortical malformation (type II lissencephaly), cerebellar abnormalities, and retinal dysplasias (Dobyns et al., 1989).

Brain malformations in dystroglycanopathies reflect the critical role that dystroglycan plays in maintaining the architecture of the neuroepithelial scaffold (Moore et al., 2002; Myshrall et al., 2012). Focal regions of retinal dysplasia have also been observed In mouse models of dystroglycanopathy, with ectopic cells protruding through the ILM (Takeda et al., 2003; Lee et al., 2005; Satz et al., 2008; Chan et al., 2010; Takahashi et al., 2011). In *Xenopus*, morpholino depletion of *dystroglycan* results in micropthalmia, degeneration of the ILM, and abnormal positioning of photoreceptors, bipolar cells, and retinal ganglion cells (Lunardi et al., 2006). However, several important questions regarding the role of dystroglycan in mammalian retinal development remain unaddressed. First, what is the underlying cause of retinal dysplasia in dystroglycanopathy? Second, is dystroglycan required for the proper migration and lamination of specific subtypes of retinal neurons? Third, how does the loss of dystroglycan affect axon guidance, dendritic stratification and mosaic spacing in the retina? Finally, how do retinal dysplasias in models of dystroglycanopathy affect the function of the retina?

Here, using multiple genetic models, we identify a critical role for dystroglycan in multiple aspects of retinal development. We show that dystroglycan within the neuroepithelium is critical for maintaining the structural integrity of the ILM. We provide *in vivo* evidence that dystroglycan’s maintenance of the ILM is required for proper neuronal migration, axon guidance, formation of synaptic lamina, and the survival of multiple retinal neuron subtypes. Surprisingly, spontaneous retinal activity appears unperturbed, despite the dramatic disruption in inner retinal development. Together, these results provide critical insight into how dystroglycan directs the proper functional assembly of retinal circuits.

## Material and Methods

### Experimental Design and Statistical Analysis

Mice were maintained on mixed genetic backgrounds and were used irrespective of sex. All phenotypic analysis was conducted with n≥3 animals obtained from at least two different litters of mice. Statistical analysis was performed using JMP Pro 13.0 software (SAS Institute). Comparison between two groups was analyzed using a Student’s t-test. Comparison between two or more groups was analyzed using a Two-Way ANOVA and Tukey post-hoc test. Comparison of retinal wave parameters was analyzed using a Wilcoxon Rank Sums test. The significance threshold was set at 0.05 for all statistical tests. * indicates p<0.05; ** indicates p<0.01; *** indicates p<0.001.

### Animals

Animal procedures were approved by OHSU Institutional Animal Care and Use Committee and conformed to the National Institutes of Health *Guide for the care and use of laboratory animals*. Animals were euthanized by administration of CO_2_. The day of vaginal plug observation was designated as embryonic day 0 (e0) and the day of birth in this study was designated as postnatal day 0 (P0). The generation and genotyping protocols for *ISPD^L79*/L79*^* (Wright et al., 2012), *DG^F/F^* (Moore et al., 2002) and *DG^β^cyt^^* (Satz et al., 2009) mice have been described previously. The presence of the cre allele in *Sox2^Cre^* (Hayashi et al., 2002), *Six3^Cre^* (Furuta et al., 2000), *Isl1^Cre^* (Yang et al., 2006) and *Nestin^Cre^* mice (Tronche et al., 1999) was detected by generic cre primers. *ISPD^+/L79*^*, *DG^β^cyt/+^^*, and *DG^F/+^*; *Six3^Cre^* or *DG^F/+^* age matched littermates were used as controls.

### Tissue Preparation and Immunohistochemistry

Embryonic retinas were left in the head and fixed overnight at 4°C in 4% PFA and washed in PBS for 30 minutes. Heads were equilibrated in 15% sucrose overnight and flash frozen in OCT medium. Postnatal retinas were dissected out of the animal and the lens was removed from the eyecup. Intact retinas were fixed at room temperature for 30 minutes in 4% PFA. Retinas were washed in PBS for 30 minutes and equilibrated in a sequential gradient of 10%, sucrose, 20% sucrose and 30% sucrose overnight. Tissue was sectioned on a cryostat at 16-25um. Tissue sections were blocked in a PBS solution containing 2% Normal Donkey Serum and 0.2% Triton for 30 minutes, and incubated in primary antibody overnight at 4°C. Sections were washed for 30 minutes and incubated in secondary antibody in a PBS solution containing 2% Normal Donkey Serum for 2-4 hours. Sections were incubated in DAPI to stain nuclei for 10 minutes, washed for 30 minutes, and mounted using Fluoromount medium. The source and concentration of all antibodies utilized in this study are listed in Table 1.

**Table 1:**
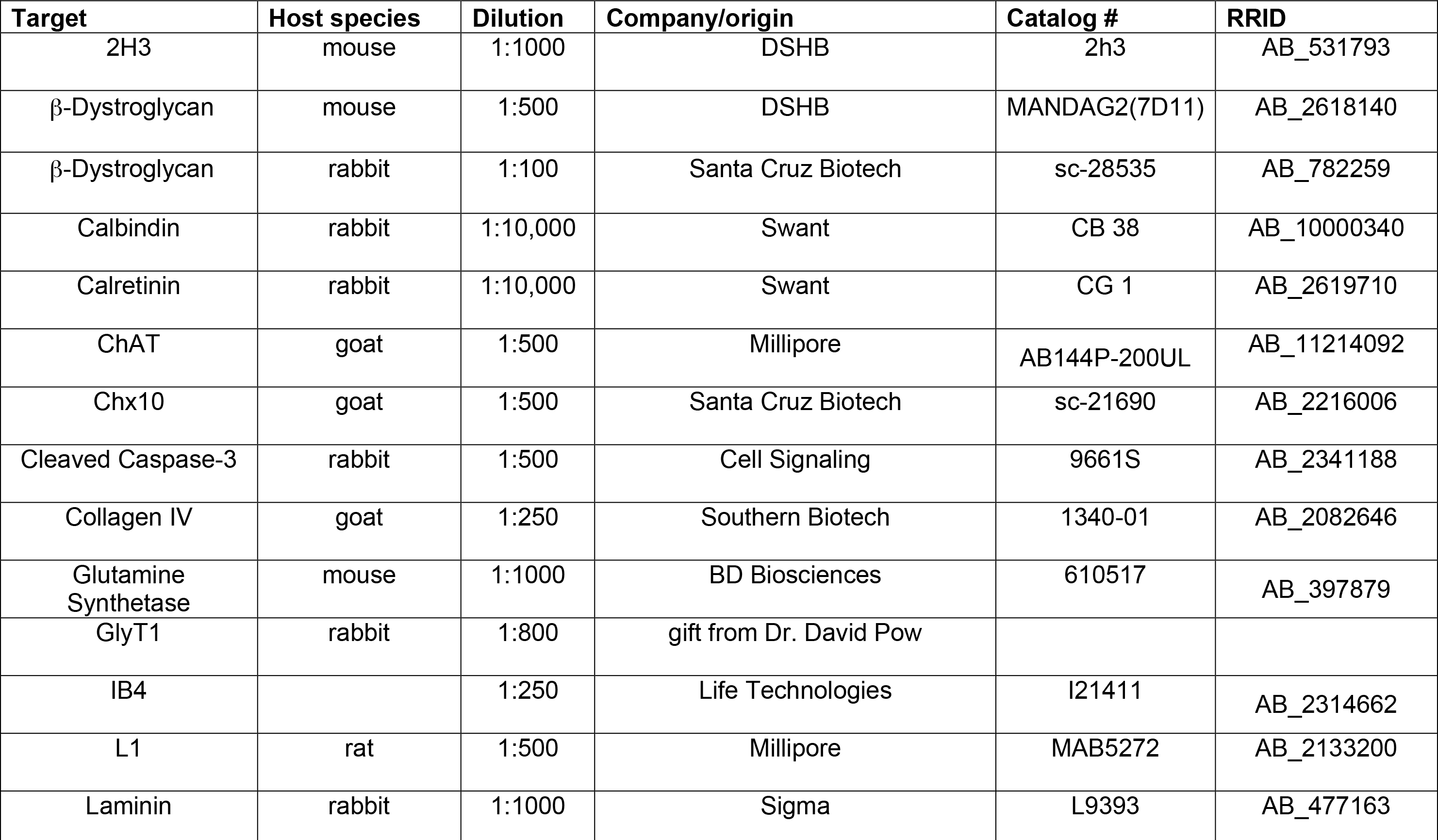
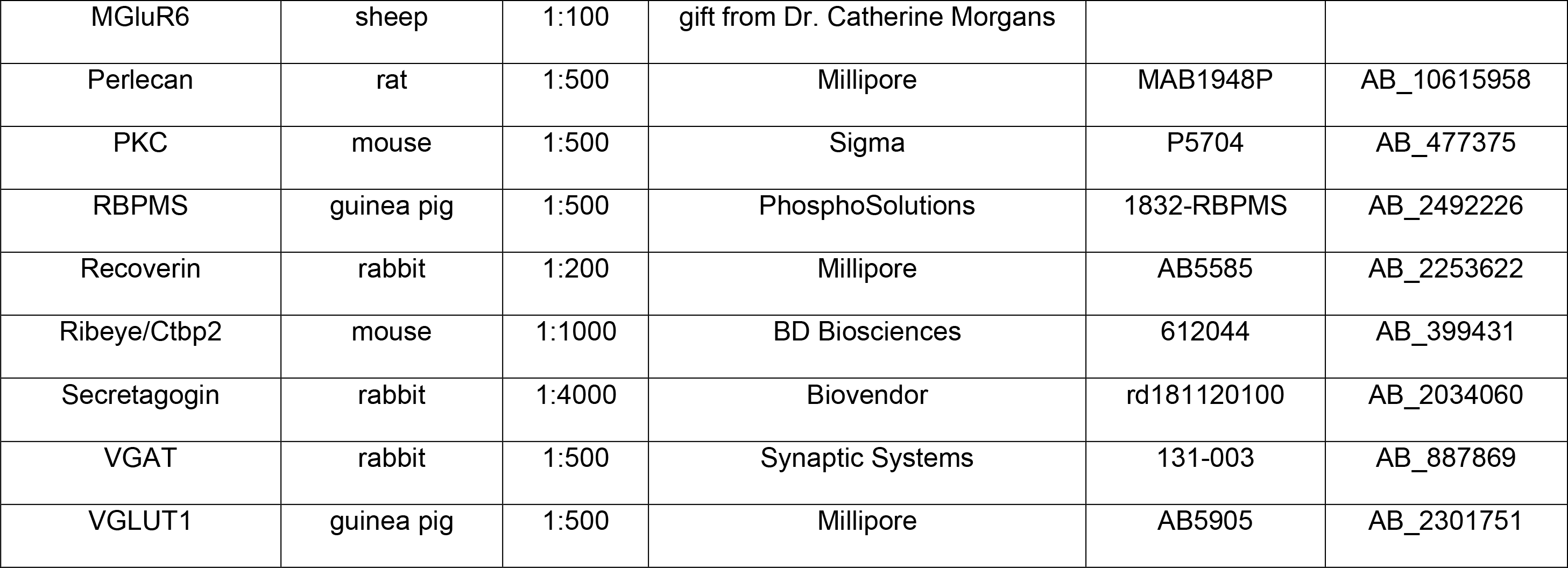
List of Antibodies. Antibodies utilized throughout the study, including target, host species, dilution, manufacturer, catalog number and RRID.

### Wholemount retinal staining

Postnatal retinas were dissected out of the animal and the lens was removed from the eyecup. Intact retinas were fixed at room temperature for 30 minutes in 4% PFA. Retinas were incubated in primary antibody diluted in PBS solution containing 2% Normal Donkey Serum and 0.2% Triton for two days at 4°C. Retinas were washed in PBS for one day and incubated in secondary antibody diluted in PBS solution containing 2% Normal Donkey Serum for two days at 4°C, washed for one day in PBS and mounted using Fluoromount medium.

### Microscopy

Imaging was performed on a Zeiss Axio Imager M2 upright microscope equipped with an ApoTome.2. Imaging of synapses was performed on a Zeiss Elyra PS.1 with LSM 710 laser-scanning confocal Super-Resolution Microscope with AiryScan. Imaging of retinal waves was performed on a Nikon TiE inverted microscope with full environmental chamber equipped with a Yokogawa CSU-W1 spinning disk confocal unit.

### Quantification of cell number and mosaic spacing

For each experiment, 3-4 locations per retina at the midpoint of each lobe were sampled. Cell counts of horizontal cells (Calbindin), apoptotic cells (Cleaved Caspase-3, P0), and starburst amacrine cells (ChAT) were obtained from 500 x 500 μm images and quantified in FIJI. Cell counts of apoptotic cells (Cleaved Caspase-3, e16) and ganglion cells (RPBMS) were obtained from 250 x 250 μm images and quantified in FIJI. Analysis of retinal mosaics (Calbindin, ChAT) were conducted on 500 x 500 μm images by measuring the X-Y coordinates for each cell and Voronoi domains were calculated in FIJI and nearest neighbor measurements calculated with WinDRP. Cell counts of proliferating cells (PH3 positive, e13, e16) were done by counting the number of positive cells in mid-retinal 20μm sections.

### Live cell imaging and analysis

Retinas from P1 *DG^F/+^*; *Six3^cre^*; *R26-LSL-GCaMP6f* and *DG^F/-^*; *Six3^cre^*; *R26-LSL-GCaMP6f* were dissected into chilled Ames’ Medium (Sigma) buffered with sodium bicarbonate and bubbled with carbogen gas (95% O_2_, 5% CO_2_). The retinas were dissected out of the eyecup, mounted RGC side up on cellulose membrane filters (Millipore) and placed in a glass-bottom petri dish containing Ames’ Medium. A platinum harp was used to stabilize the filter paper during imaging. Imaging was performed at 30°C using a 10x0.45 Plan Apo Air objective with a field of 1664 by 1404 μm with a 3Hz imaging timeframe. The field was illuminated with a 488 nm laser. 3-4 retinal fields were imaged per retina, and each field of retina was imaged for a two-minute time series using a 300ms exposure and each field was sampled 3-5 times per imaging session.

Thirty representative control and thirty representative mutant time series were randomly selected for analysis. Only waves that initiated and terminated within the imaging field were used for analysis. To measure wave area, movies were manually viewed using FIJI frame by frame to determine the start and end frame of a wave. A Z-Projection for maximum intensity was used to create an image with the entire wave, and the boundary of the wave was manually traced to determine the area. Wave area per time was calculated by dividing the area of the wave by the duration in seconds of the wave. Any wave lasting less than 2 seconds was not used in analysis, consistent with previous studies (Blankenship et al., 2009).

## Results

### Dystroglycan is required for inner limiting membrane integrity

Dystroglycan plays a critical role in the developing cortex, where it anchors radial neuroepithelial endfeet to the basement membrane along the pial surface. In the absence of functional dystroglycan, disruptions in the cortical basement membrane and detachment of neuroepithelial endfeet lead to profound neuronal migration phenotypes (Moore et al., 2002; Myshrall et al., 2012). In the adult retina, dystroglycan is present in blood vessels, in RGCs, at ribbon synapses in the OPL, and at the ILM, which serves as a basement membrane that separates the neural retina from the vitreous space (Montanaro et al., 1995; Omori et al., 2012). However, the role of dystroglycan in regulating neuronal migration, axon guidance or dendritic stratification of specific cell types during retinal development has not been examined in a comprehensive manner. To address this open question, we first examined the expression pattern of dystroglycan in the developing retina. Using immunohistochemistry, we observed dystroglycan expression along radial processes that span the width of the retina, and its selective enrichment at the ILM from embryonic ages (e13) through birth (P0) (Figure 1A). These processes are likely a combination of neuroepithelial cells and the basal process of migrating RGCs. Loss of staining in retinas from an epiblast-specific *dystroglycan* conditional knockout (*DG^F/-^*; *Sox2^cre^*) confirmed the specificity of this expression pattern (Figure 1B).

**Figure 1:**
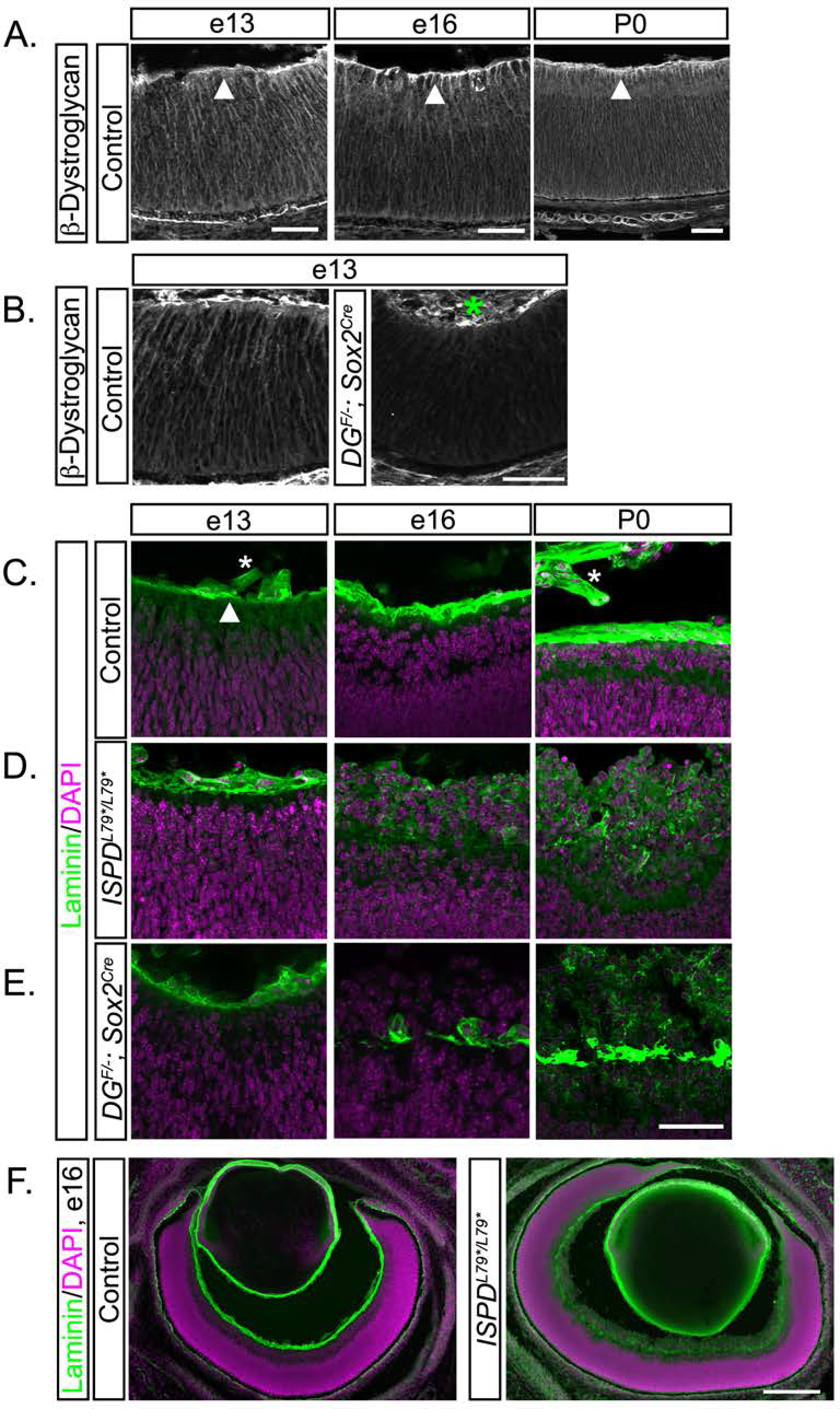
The inner limiting membrane undergoes progressive degeneration in the absence of functional dystroglycan. (A) Dystroglycan (β-DG) is expressed throughout the developing retina, with an enrichment at the inner limiting membrane (ILM). (B) Dystroglycan expression is lost in *DG^F/-^*; *Sox2^Cre^* mice. (C-E) The initial assembly of the ILM (laminin, green) occurs normally in the absence of functional dystroglycan in (D) *ISPD^L79*/L79*^* and (E) *DG^F/-^*; *Sox2^cre^* retinas. The ILM in *ISPD^L79*/L79*^* and *DG^F/-^*; *Sox2^cre^* retinas undergoes progressive degeneration at e16 (middle) and P0 (right), and retinal neurons migrate into the vitreous. (F) The ILM in *ISPD^L79*/L79*^* retinas undergoes degeneration (right panel) that is present across the entire span of the retina at e16. Arrowheads indicate ILM, asterisks indicate non-specific staining of blood vessels. Scale bar, 50μm A-D, 200 μm E.

The first step in the assembly of basement membranes is the recruitment of laminin polymers to the cell surface by sulfated glycolipids, followed by the stabilization of laminin polymers by transmembrane receptors (Yurchenco, 2011). To determine whether dystroglycan is required for the initial formation of the ILM during retinal development, we utilized two complementary genetic models. *ISPD^L79*/L79*^* mutants, previously identified in a forward genetic screen, lack the mature glycan chains required for dystroglycan to bind ligands such as laminin and are a model for severe dystroglycanopathy (Wright et al., 2012). *DG^F/-^*; *Sox2^cre^* conditional mutants lack dystroglycan in epiblast-derived tissues, including all retinal tissue, and were utilized to confirm that phenotypes observed in *ISPD^L79*/L79*^* mice are dystroglycan dependent. The enrichment of laminin at the ILM appeared normal in early retinal development at e13 in both *ISPD^L79*/L79*^* and *DG^F/-^*; *Sox2^cre^* mutants (Figure 1C-E), indicating that dystroglycan is not required for the initial formation of the ILM. However, at e16, we observed a loss of laminin staining and degeneration of the ILM across the entire surface of the retina in *ISPD^L79*/L79*^* and *DG^F/-^*; *Sox2^cre^* mutants (Figure 1D-F). The loss of ILM integrity in *ISPD^L79*/L79*^* and *DG^F/-^*; *Sox2^cre^* mutants was accompanied by the inappropriate migration of retinal neurons, resulting in the formation of an ectopic layer of neurons protruding into the vitreous space.

Following the initial polymerization of laminin on cell surfaces, additional ECM proteins bind and crosslink the nascent basement membrane to increase its stability and complexity. Examination of *ISPD^L79*/L79*^* and *DG^F/-^*; *Sox2^cre^* mutant retinas at P0 revealed a loss of the ECM proteins Collagen IV (red), and Perlecan (green), coinciding with the disruptions in Laminin (purple) (Figure 2A, B, C). These data suggest that while dystroglycan is not required for the initial formation of the ILM, it is essential for its maturation and maintenance. Furthermore, dystroglycan is critical for the ILM to function as a structural barrier to prevent the ectopic migration of neurons into the vitreous space.

**Figure 2:**
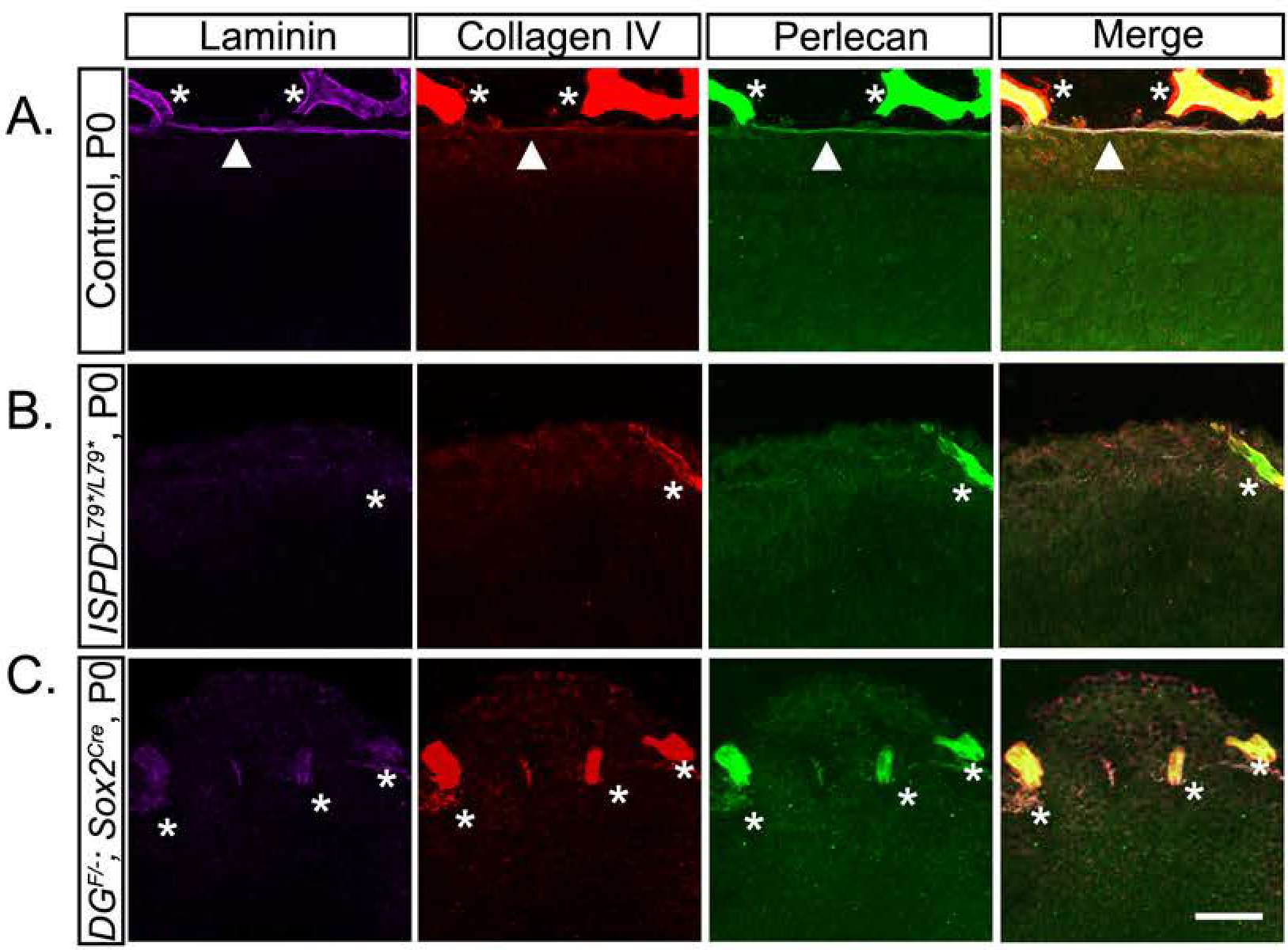
Dystroglycan is required to localize ECM proteins at the ILM. (A) Multiple extracellular matrix proteins, including laminin (purple), collagen IV (red), and perlecan (green), localize to the ILM in controls. (B**-C**) Localization of extracellular matrix proteins is disrupted in the ILM in the absence of functional dystroglycan in *ISPD^L79*/L79*^* (B) and *DG^F/-^*; *Sox2^cre^* (C) retinas at P0. Arrowheads indicate ILM, asterisks indicate blood vessels. Scale bar, 50μm.

### Dystroglycan is required for vascular and optic fiber layer development

The hyaloid vasculature in the embryonic retina normally regresses as astrocytes and the retinal vascular plexus emerge through the optic nerve head beginning around birth (Fruttiger, 2007). Previous studies have found that defects in ILM integrity disrupt the emergence and migration of astrocytes and the retinal vasculature (Lee et al., 2005; Edwards et al., 2010; Takahashi et al., 2011; Tao and Zhang, 2016). In agreement with these findings, we observe that at embryonic ages in *ISPD^L79*/L79*^* mutant retinas, the hyaloid vasculature becomes embedded within the ectopic retinal neuron layer at e16 (Figure 3A), and fails to regress at P0 (Figure 3B). In addition, the emergence of the retinal vasculature and astrocytes is stunted in *ISPD^L79*/L79*^* mutants (Figure 3B).

**Figure 3:**
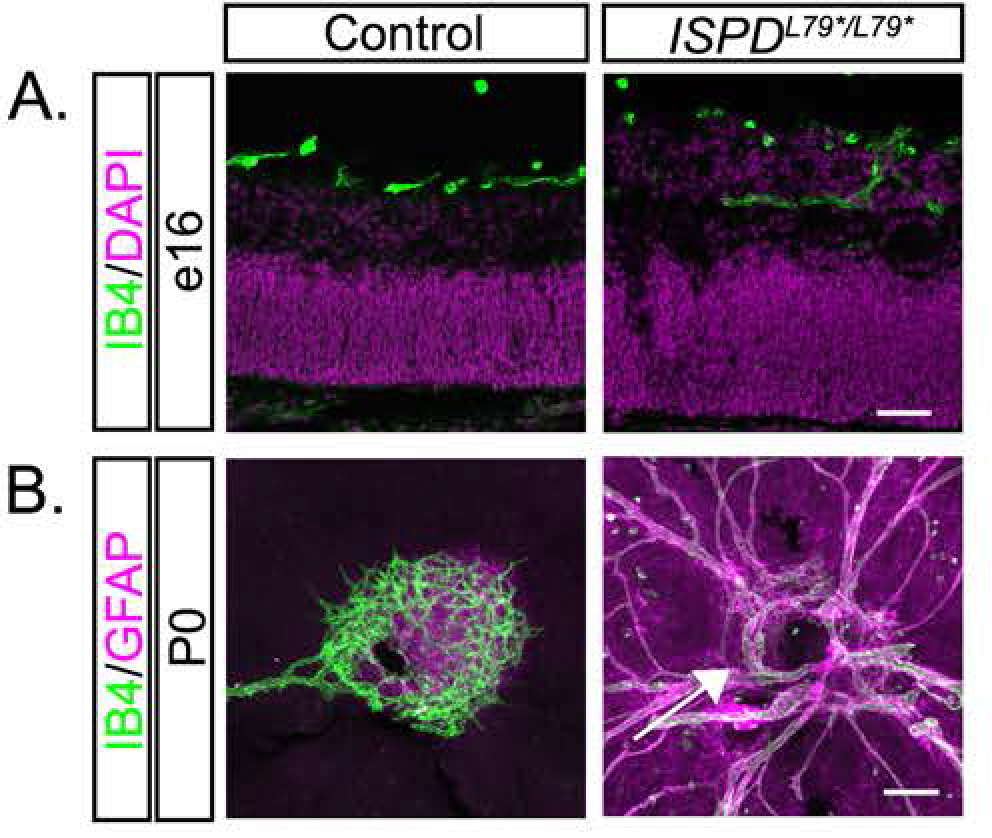
Dystroglycan regulates normal retinal vasculature development. (A) IB4 labeled hyaloid vasculature is present in the vitreous adjacent to the GCL in control retinas (left) but is embedded within ectopic cell clusters *ISPD^L79*/L79*^* (right) retinas at e16. (B) Flat mount retinas at P0 show the emergence of the primary vascular plexus (IB4, green) and astrocytes (GFAP, purple) in controls (left). In *ISPD^L79*/L79*^* retinas (right), the emergence of astrocytes and the primary vascular plexus into the retina is delayed (arrow) and there is a persistence of hyaloid vasculature. Scale bar 50 μm A, 100 μm B.

We have shown previously that the organization of the basement membrane by dystroglycan provides a permissive growth substrate for axons in the developing spinal cord (Wright et al., 2012). In addition, contact with laminin in the ILM stabilizes the leading process of newly generated RGCs to direct the formation of the nascent axon (Randlett et al., 2011). These axons remain in close proximity to the ILM as they extend centrally towards the optic nerve head, forming the optic fiber layer (OFL). Therefore, we examined whether the disruptions in the ILM affected the guidance of RGC axons in *ISPD^L79*/L79*^* and *DG^F/-^*; *Sox2^cre^* mutants. At e13, axons in both control and mutant retinas formed a dense and continuous network in the basal retina, directly abutting the ILM (Figure 4A). In contrast, at e16 (Figure 4B) and P0 (Figure 4C), RGC axons in *ISPD^L79*/L79*^* and *DG^F/-^*; *Sox2^cre^* mutants were disorganized, exhibiting both defasciculation (asterisks) and hyperfasciculation (arrowheads).

**Figure 4:**
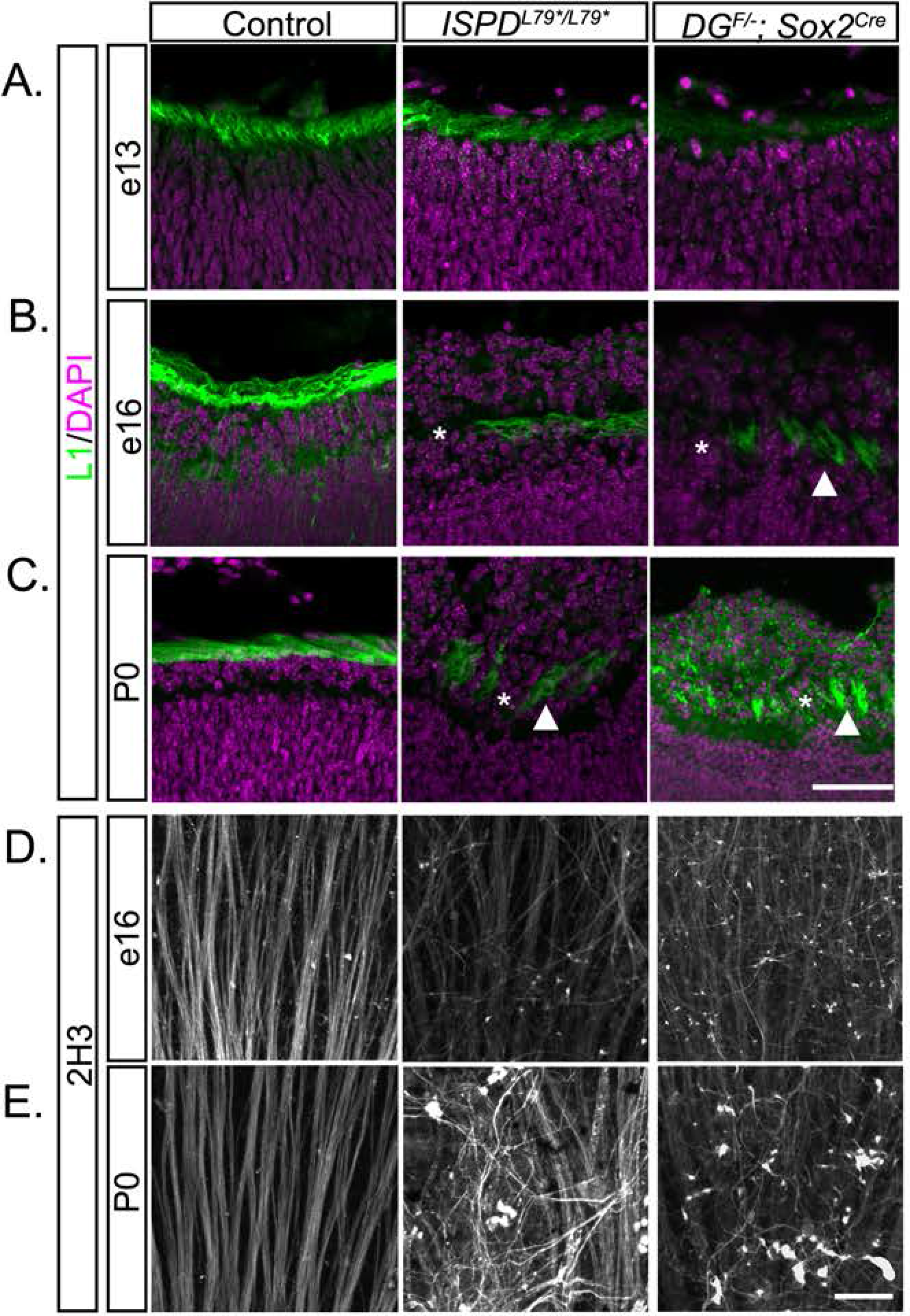
Dystroglycan is required for intraretinal axon guidance, (A) L1 positive axons in the optic fiber layer (OFL) initially appear normal in *ISPD^L79*/L79*^* (middle), and *DG^F/-^*; *Sox2^cre^* retinas (right) at e13. (B, C) As the ILM degenerates in *ISPD^L79*/L79*^* and *DG^F/-^*; *Sox2^cre^* retinas at e16 (B) and P0 (C), axons hyperfasciculate (arrowhead) and exhibit defasciculation (asterisk) within the OFL. (D, E). Flat mount preparations from *ISPD^L79*/L79*^* and *DG^F/-^*; *Sox2^cre^* retinas at e16 (D) and P0 (E) show progressive disruption of axon tracts (Neurofilament, 2H3). Scale bar, 50 μm.

To gain further insight into the specific defects that occur in RGC axons, we used a flat mount retina preparation. Regardless of their location in the retina, all RGCs orient and extend their axons towards the center of the retina, where they exit the retina through the optic nerve head (Bao, 2008). In control retinas at e16 (Figure 4D, left panel) and P0 (Figure 4E, left panel) axons traveled towards the optic nerve head in fasciculated, non-overlapping bundles. In contrast, we frequently observed defasciculated RGC axons that grew in random directions without respect to their orientation to the optic nerve head in both *ISPD^L79*/L79*^* and *DG^F/-^*; *Sox2^cre^* mutants. Together, these data show that proper growth and guidance of RGC axons to the optic nerve head requires dystroglycan to maintain an intact ILM as a growth substrate.

### Dystroglycan is required for axonal targeting, dendritic lamination, and cell spacing in the postnatal retina

Our results in *ISPD^L79*/L79*^* and *DG^F/-^*; *Sox2^cre^* mutants demonstrate that dystroglycan is required for ILM integrity and to prevent the ectopic migration of neurons into the vitreous (Figure 1). However, the specific neuronal subtypes affected in models of dystroglycanopathy are unknown, and the role of dystroglycan in regulating postnatal aspects of retinal development has not been examined. The synaptic layers of the retina develop postnatally, with tripartite synapses between the photoreceptors, bipolar cells and horizontal cells forming in the OPL, and synapses between bipolar cells, amacrine cells and retinal ganglion cells forming in the IPL. The development of these synaptic layers requires the precise stratification of both axons and dendrites that occurs between P0 and P14. Since *ISPD^L79*/L79*^* and *DG^F/-^*; *Sox2^cre^* mutant mice exhibit perinatal lethality, we deleted *dystroglycan* selectively from the early neural retina using a *Six3^Cre^* driver (Furuta et al., 2000) (Figure 5A).

**Figure 5:**
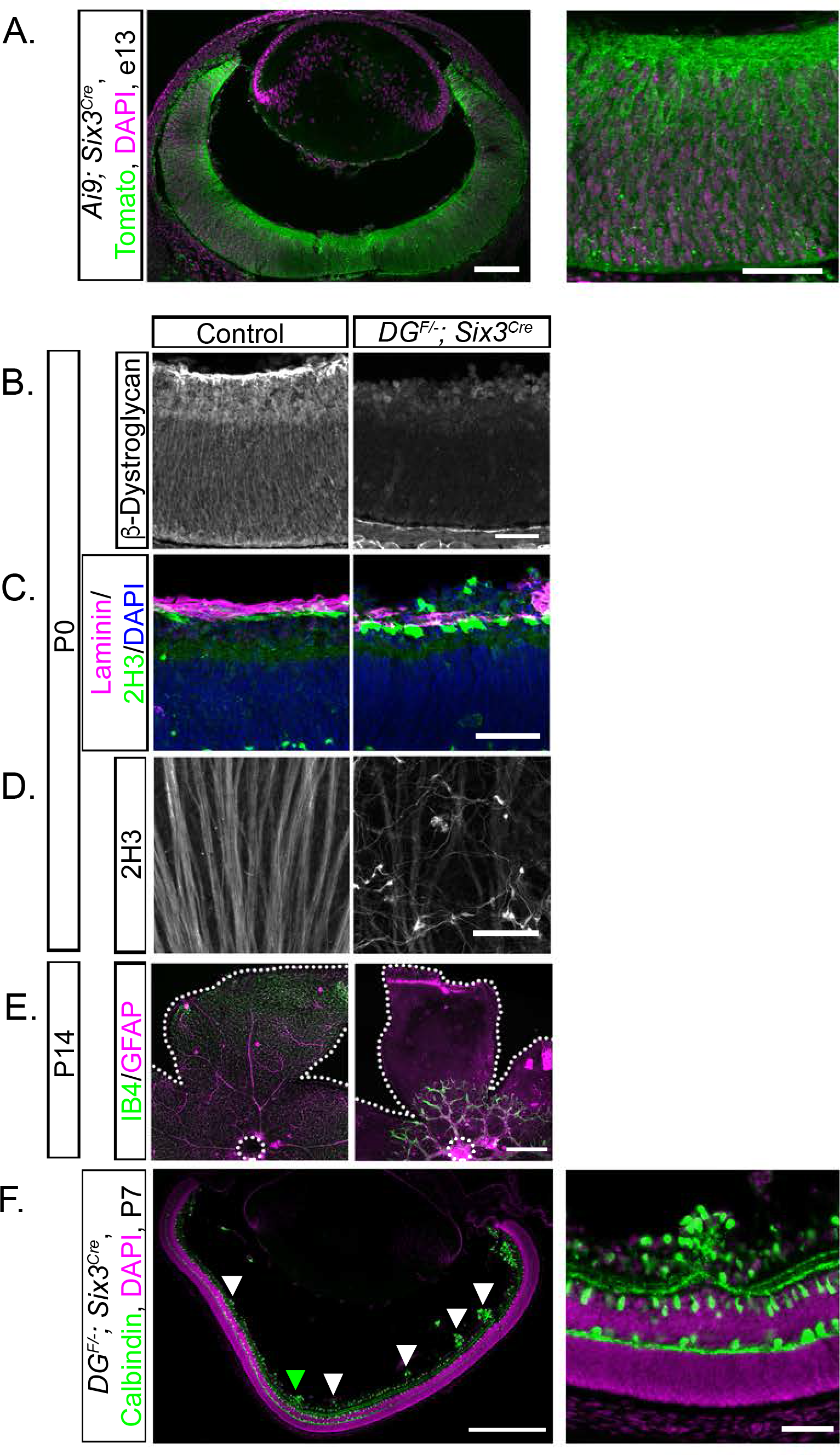
Conditional deletion of *dystroglycan* in the developing retina results in migration and axon guidance defects. (A) Recombination pattern of *Rosa26-lox-stop-lox-TdTomato; Ai9* reporter (green) by *Six3^cre^* shows expression throughout the retina and in axons at e13. (B) Dystroglycan protein expression is lost in *DG^F/-^*; *Six3^cre^* mice. (C, D) *DG^F/-^*; *Six3^cre^* (right) mice exhibit inner limiting membrane degeneration (top, purple, laminin) and abnormal axonal fasciculation and guidance (top, green, bottom, 2H3). (E) Primary vascular plexus (IB4, green) and astrocytes (GFAP, purple) migrate from the optic nerve head (dashed circle) to the edge of the retina (dashed line) in control retinas at P14 (right). Vascular and astrocyte migration is stunted in *DG^F/-^*; *Six3^cre^* retinas. (F) Focal migration defects (arrowheads) in P7 *DG^F/-^*; *Six3^cre^* retinas are present across the entire span of the retina. Green arrowhead indicates high magnification image in F (right). Scale bar 100 μm A left, 500 μm F left, E, 50 μm A, D right, B-D.

Analysis of *DG^F/-^*; *Six3^cre^* retinas confirmed that dystroglycan protein was lost along the neuroepithelial processes and at the ILM (Figure 5B). We next confirmed that *DG^F/-^*; *Six3^cre^* mice recapitulated the retinal phenotypes identified in *ISPD^L79*/L79*^* and *DG^F/-^*; *Sox2^cre^* mice. We observed a degeneration of the ILM (laminin, purple) accompanied by ectopic migration of neurons into the vitreous (Figure 5C), abnormal fasciculation and guidance of RGC axons (Figure 5D), and defective emergence and migration of astrocytes and the vascular plexus (Figure 5E). While fully penetrant, the ILM degeneration and neuronal migration defects in *DG^F/-^*; *Six3^cre^* mice were milder than in *DG^F/-^*; *Sox2^cre^* mice, exhibiting a patchiness that was distributed across the retina (Figure 5F). The defects in *DG^F/-^*; *Six3^cre^* mice contrast the finding that conditional deletion of *dystroglycan* with *Nestin^cre^* does not affect the overall structure of the retina (Satz et al., 2009). We find that this difference is likely due to the onset and pattern of *Cre* expression, as recombination of a *Cre*-dependent reporter (*Rosa26-lox-stop-lox-TdTomato; Ai9*) occurred earlier and more broadly in *Six3^Cre^* mice than in *Nestin^cre^* mice (Figure 5A, data not shown).

*DG^F/-^*; *Six3^cre^* mice are healthy and survive into adulthood, allowing us to examine the role of dystroglycan in postnatal retinal development. We analyzed *DG; Six3^cre^* retinas at P14 when migration is complete and the laminar specificity of axons and dendrites has been established (Morgan and Wong, 1995). The overall architecture of the ONL appeared unaffected by the loss of dystroglycan in *DG^F/-^*; *Six3^cre^* mice, and cell body positioning of photoreceptors appeared similar to controls (Figure 6A). Within the INL, the laminar positioning of rod bipolar cell bodies (PKC, Figure 6B), cone bipolar cell bodies (SCGN, Figure 6C), horizontal cells (arrows, Calbindin, Figure 6E), and Müller glia cell bodies (Figure 6H) and the targeting of their processes to the OPL appeared normal in *DG^F/-^*; *Six3^cre^* mutants. However, bipolar cell axons that are normally confined to the synaptic layers in the IPL, and Müller glia processes that are normally concentrated at the ILM both extended aberrant projections into the ectopic clusters (Figure 6B-C, H).

**Figure 6:**
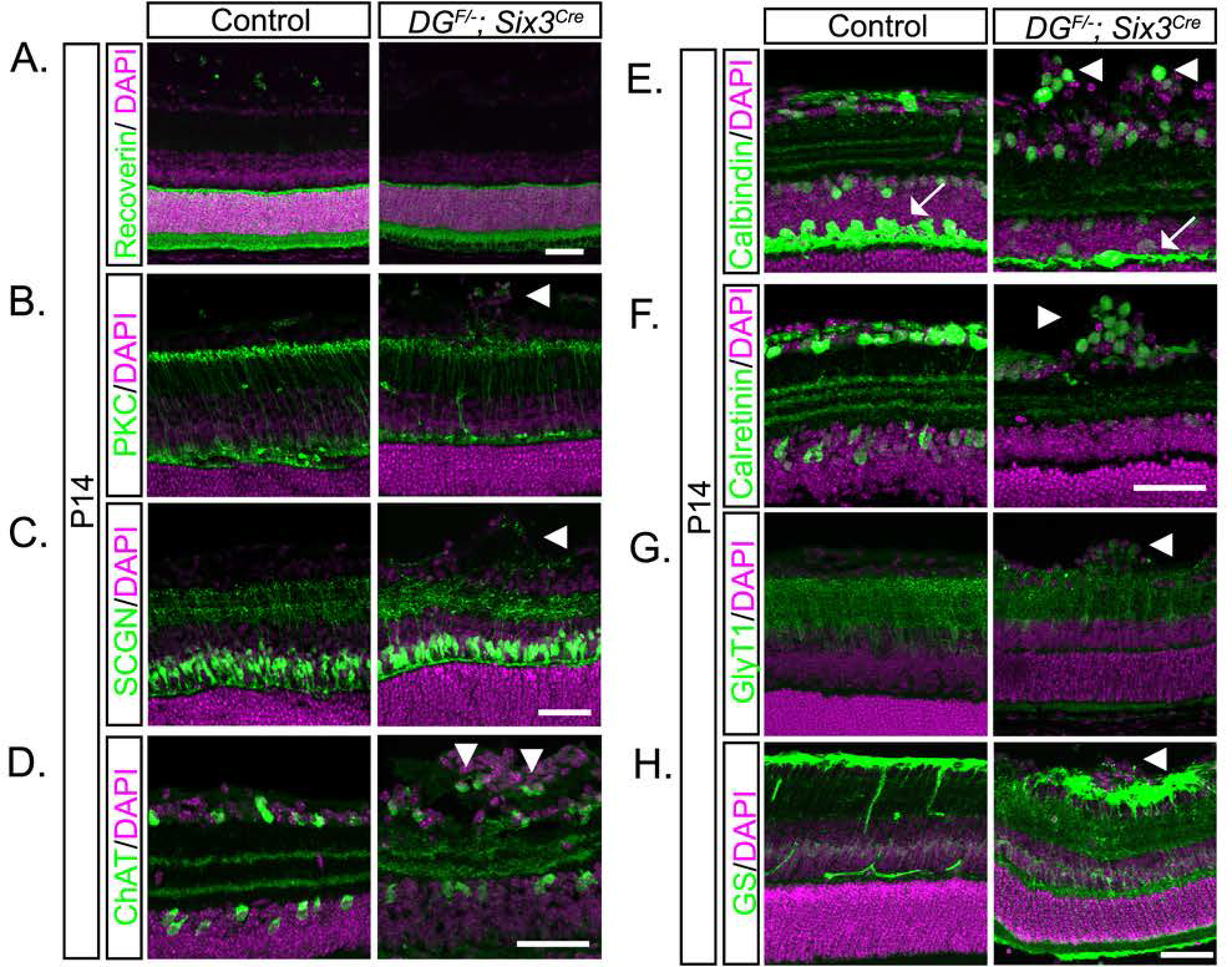
Disrupted postnatal circuit formation in the inner retina of *dystroglycan* mutants. (A) Photoreceptors (recoverin) have normal lamination in P14 *DG^F/-^*; *Six3^cre^* retinas (right). (B-C) The cell bodies of bipolar cells (PKC, B, SCGN, C) exhibit normal lamination patters, while their axons extend into ectopic cellular clusters in the ganglion cell layer. (D-G) Abnormal cellular lamination and disruptions in dendritic stratification of multiple amacrine and retinal ganglion cell types is observed in *DG^F/-^*; *Six3^cre^* retinas. (D) ChAT labels starburst amacrine cells, (E) calbindin and (F) calretinin label amacrine and ganglion cells, and (G) GlyT1 labels glycinergic amacrine cells. (H) Müller glia (glutamine synthetase) cell bodies are normally positioned while their inner retinal processes extend into ectopic cellular clusters. Arrowheads indicate axons or cell bodies in ectopic clusters, arrows indicate horizontal cell layer. Scale bar, 50 μm.

In contrast to the normal laminar architecture of the outer retina, the inner retina was disorganized in *DG^F/-^*; *Six3^cre^* mutants. Subsets of amacrine and ganglion cells labeled by ChAT (starburst amacrine cells, Figure 6D), calbindin (Figure 6E), and calretinin (Figure 6F) that are normally confined to the INL and GCL were present in the ectopic clusters that protrude into the vitreous space. Glycinergic amacrine cells (GlyT1, Figure 6G), whose cell bodies are normally found in a single layer within the INL, were also present within ectopic clusters. Compared to the OPL, which appeared grossly normal, dendritic stratification within the IPL in *DG^F/-^*; *Six3^cre^* mutant retinas was disrupted. The laminated dendritic strata appeared expanded (Figure 6D), fragmented (Figure 6D-F) and occasionally lacked an entire lamina (Figure 6E). These defects were restricted to regions of the retina where ectopic neuronal clusters were present, whereas regions of the *DG^F/-^*; *Six3^cre^* mutant retina with normal cellular migration and lamination also had normal dendritic stratification (Figure 5F). These results demonstrate that the ectopic clusters consisted of multiple subtypes of amacrine cells and ganglion cells that normally reside in the INL and GCL, and that the disorganization of the dendritic strata are likely secondary to the cell migration defects.

Over the course of retinal development, multiple cell types, including horizontal cells and amacrine cells, develop mosaic spacing patterns that ensure cells maintain complete and non-random coverage over the surface of the retina (Wassle and Riemann, 1978). This final mosaic pattern is established by both the removal of excess cells through apoptosis and the lateral dispersion of “like-subtype” cells via homotypic avoidance mechanisms (Kay et al., 2012; Li et al., 2015). To determine whether the defects in establishing proper laminar positioning of retinal subtypes in *DG^F/-^*; *Six3^cre^* mutants extends to mosaic spacing, we performed nearest neighbor analysis. For horizontal cells, which exhibit normal lamination in *DG^F/-^*; *Six3^cre^* mutant retinas, we observed a small, but significant reduction in the number of cells (Figure 7A-B, Two-Way ANOVA, p<0.0001, Tukey HSD post-hoc test ***p<0.0001). Despite the reduction in horizontal cell number, nearest neighbor curves between controls and mutants are the same shape, indicating that horizontal cell mosaics are maintained in dystroglycan mutants (Figure 7A-B).

**Figure 7:**
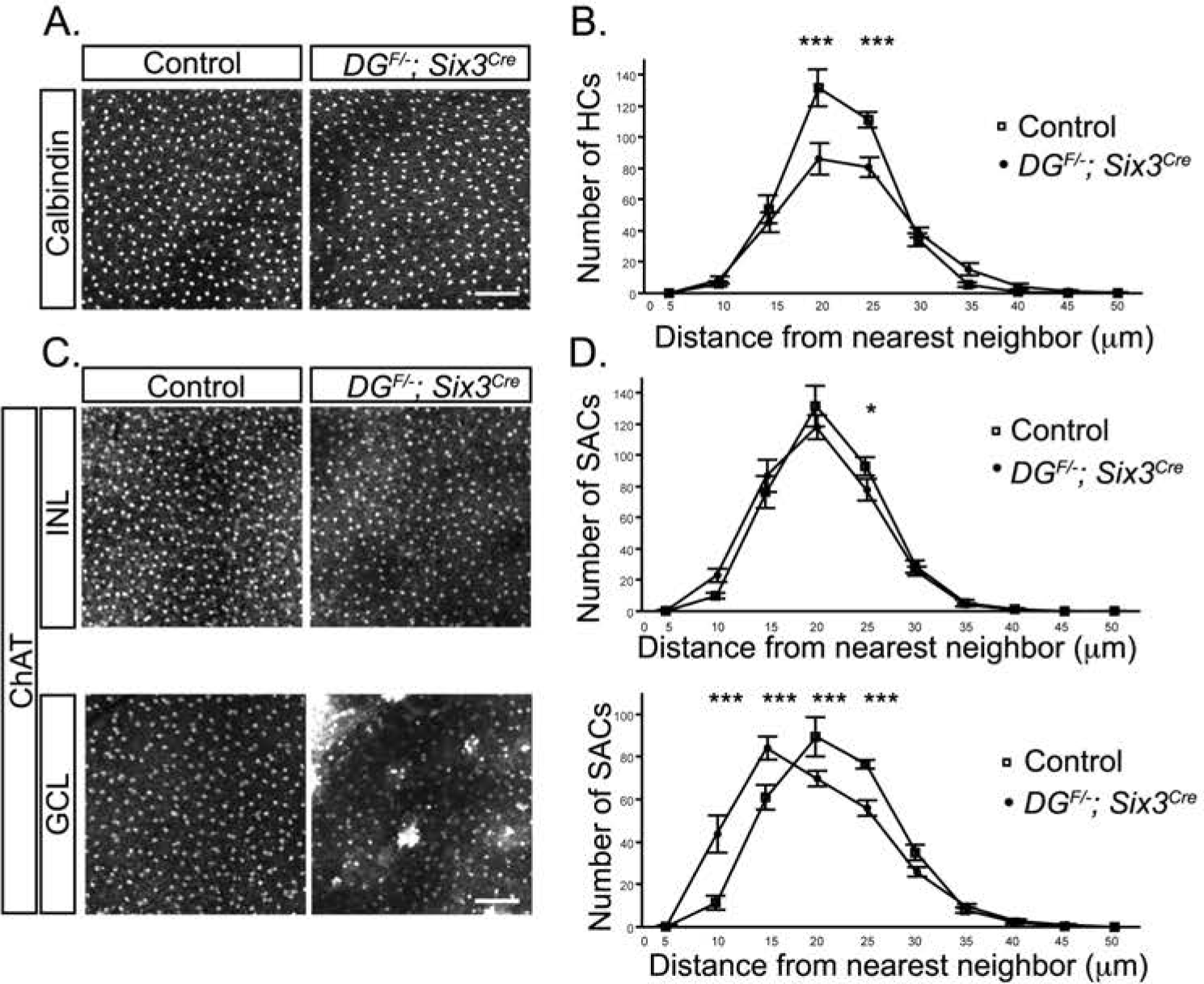
Dystroglycan is required for mosaic cell spacing in the ganglion cell layer. (A, B). Horizontal cells (2H3) in flat mount P14 adult retinas have reduced cellular density, but normal mosaic cell spacing curves (Nearest neighbor analysis, p<0.0001, Two-Way ANOVA, Tukey HSD post-hoc test ***p<0.0001, n=20 samples from 5 control retinas, 18 samples from 5 mutant retinas). (C, D) Mosaic cell spacing of starburst amacrine cells (ChAT) in the inner nuclear layer is normal (Nearest neighbor analysis, Two-Way ANOVA, Tukey HSD post-hoc test *p=0.0494, n=10 samples from 3 control retinas, 10 samples from 3 mutant retinas), while the ectopic clustering of starburst amacrine cells in the ganglion cell layer results in a significant disruption of mosaic spacing (Nearest neighbor analysis, p<0.0001, Two-Way ANOVA, Tukey HSD post-hoc test ***p<0.0001, n=10 samples from 3 control retinas, 10 samples from 3 mutant retinas). Scale bar 100μm.

ChAT positive starburst amacrine cells are present in two distinct lamina that form mosaic spacing patterns independent from one another. Consistent with this, ChAT labeled cells in the INL showed normal mosaic cell spacing (Figure 7C, top, Figure 7D, top, Two-Way ANOVA, Tukey HSD post-hoc test *p=0.0494). In contrast, the GCL contained prominent ChAT positive clusters that corresponded to the ectopic protrusions that extend into the vitreous, resulting in a decrease in cell spacing as determined by nearest neighbor analysis (Figure 7C, bottom, Figure 7D, bottom, Two-Way ANOVA, p<0.0001, Tukey HSD post-hoc test ***p<0.0001). These results demonstrate that laminar migration defects in *DG^F/-^*; *Six3^cre^* mutants degrade the mosaic spacing of cells in the GCL, and that contact with an intact ILM is likely required for the proper lateral dispersion of these cells.

### Deletion of dystroglycan leads to a loss of photoreceptors, horizontal cells and ganglion cells

During development, normal physiological apoptotic cell death during the first two postnatal weeks plays an important role in retinal maturation (Young, 1984). This process is critical for establishing the proper numbers and spacing of some subtypes of cells across the mature retina, as well as removing cells that fail to connect to appropriate synaptic targets (Braunger et al., 2014). Degeneration of the ILM during development can lead to a reduction in the number of ganglion cells, and previous analysis of dystroglycanopathy mutants has noted thinning of the retina (Halfter et al., 2005; Lee et al., 2005; Satz et al., 2008; Chan et al., 2010; Takahashi et al., 2011). In agreement with these results, we observed that the retinas of *DG^F/-^*; *Six3^cre^* mutants are thinner (Figure 6, 8). However, the specific cell types affected by the loss of dystroglycan in the retina are unknown.

**Figure 8:**
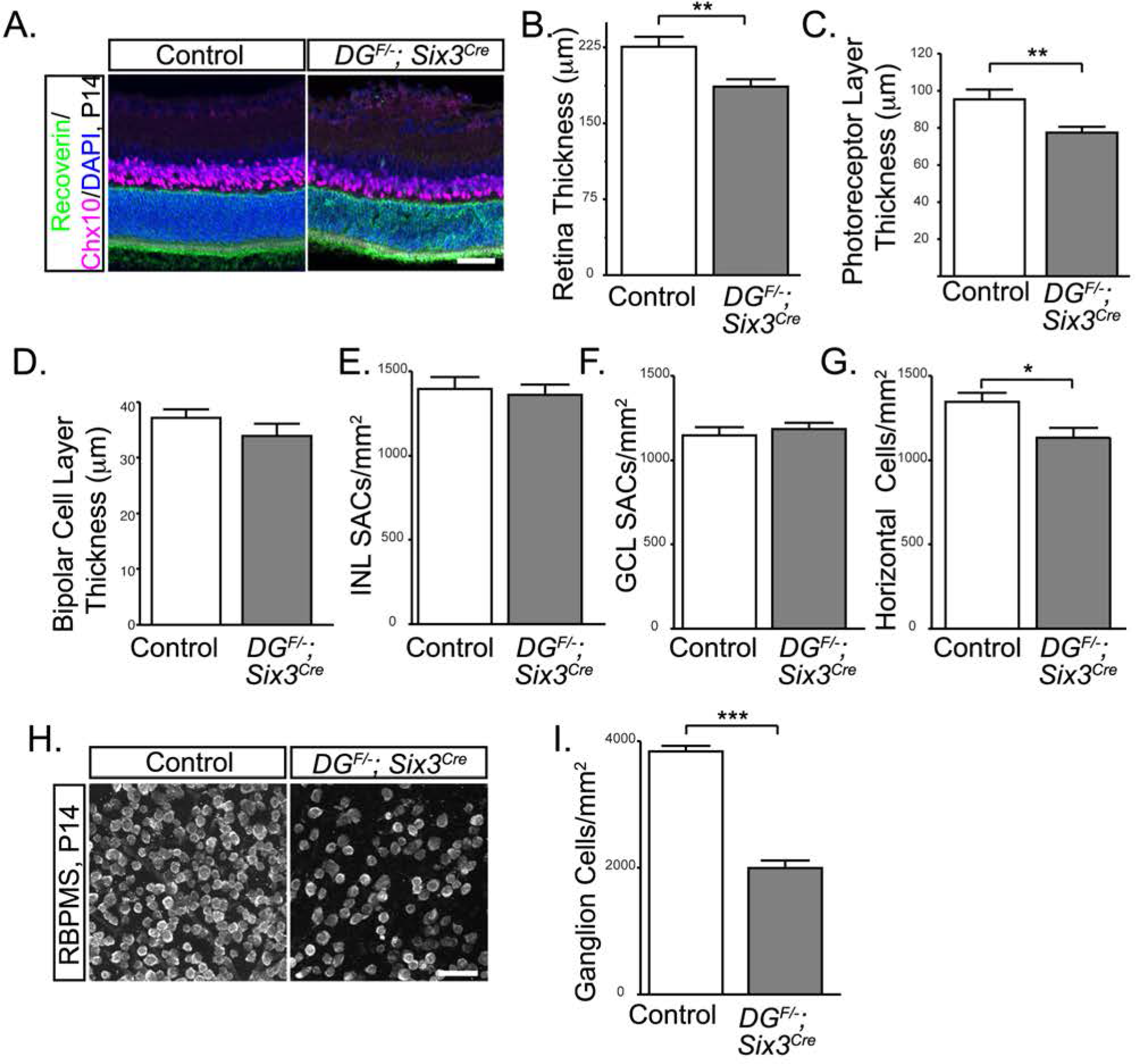
Retinal thinning in *dystroglycan* mutants. (A-D) P14 *DG^F/-^*; *Six3^cre^* retinas show decreased retinal thickness (DAPI, p=0.0039, t test, n=7 control, 7 mutant), a decreased thickness of the photoreceptor layer (C, Recoverin, green, p=0.0163, t-test, n=7 control, 7 mutant) and no change in thickness of the bipolar cell layer (D, Chx10, purple, p>0.05, t-test, n=7 control, 7 mutant). Starburst amacrine cell density is normal in both the INL (E, p>0.05, t-test, n=10 samples from 3 control retinas, 10 samples from 3 mutant retinas)and GCL (F, p>0.05, t-test, n=10 samples from 3 control retinas, 10 samples from 3 mutant retinas), while horizontal cells show a slight reduction in cell density (G) in P14 *DG^F/-^*; *Six3^cre^* retinas (p=0.0112, t test, n=20 samples from 5 control retinas, 18 samples from 5 mutant retinas). (H, I) Ganglion cell density (RBPMS) is reduced by approximately 50% in P14 *DG^F/-^*; *Six3^cre^* retinas (p<0.0001, t-test, n=12 samples from 3 control retinas, 12 samples from 3 mutant retinas). Scale bar 50 μm.

To investigate the mechanism by which loss of dystroglycan contributes to retinal thinning, we began by measuring the distance between the edges of the inner and outer retina in control and *DG^F/-^*; *Six3^cre^* mutants by DAPI staining and found that there was a significant reduction in overall retinal thickness by approximately 20% in mutants (Figure 8A, blue, Figure 8B, t test, p=0.0039). We next investigated which specific cell types contribute to retinal thinning. In the outer retina, the thickness of the photoreceptor layer (recoverin, Figure 8A, green, Figure 8C, t-test, p=0.0163) was reduced by approximately 20% and the density of horizontal cells had a small, yet significant reduction in *DG^F/-^*; *Six3^cre^* mutants (calbindin, Figure 8G, t-test, p=0.0112). In contrast, the thickness of the bipolar cell layer (Chx10, Figure 8A, purple, Figure 8D, t-test, p>0.05) was normal. In the inner retina, there was a 50% reduction in the density of ganglion cells (Figure 8H, I, t-test, p<0.0001), while the density of ChAT+ starburst amacrine cells in both the INL and GCL was normal in *DG^F/-^*; *Six3^cre^* mutants (Figure 8E, F, t-test, p>0.05). Thus, a reduction in the number of photoreceptors, horizontal cells and RGCs contribute to the overall thinning of *DG^F/-^*; *Six3^cre^* retinas.

To determine whether the reduction in photoreceptors, horizontal cells and RGCs in the absence of dystroglycan was due to defects in proliferation of retinal progenitors, we examined phospho-Histone H3 (PH3) staining at embryonic ages. PH3 positive mitotic progenitors were localized along the apical surface of the retina, and were present at the normal number in *ISPD^L79*/L79*^* mutants at e13 and e16 and (Figure 9 A, B, t-test, p>0.05). To determine whether the reduced number of neurons in mutant retinas was due to increased apoptosis, we quantified the number of caspase-3 positive cells. At e13, we observed no difference between in the number of caspase-3 positive cells in *ISPD^L79*/L79*^* mutants and controls (data not shown). In contrast, there was a significant increase in caspase-3 positive cells in *ISPD^L79*/L79*^* mutants at e16 (Figure 9D, t-test, p<0.0001) and P0 (Figure 9C, D, t-test, p<0.0001) that was restricted to the ganglion cell layer. Similarly, we observed an increased number of cleaved caspase-3 positive cells in the ganglion cell layer in *DG^F/-^*; *Six3^cre^* mutants at P0 (Figure 9C, D, t-test, p=0.0279). These results led us to conclude that the loss of RGCs in *DG^F/-^*; *Six3^cre^* mutants is due to increased apoptotic cell death.

**Figure 9:**
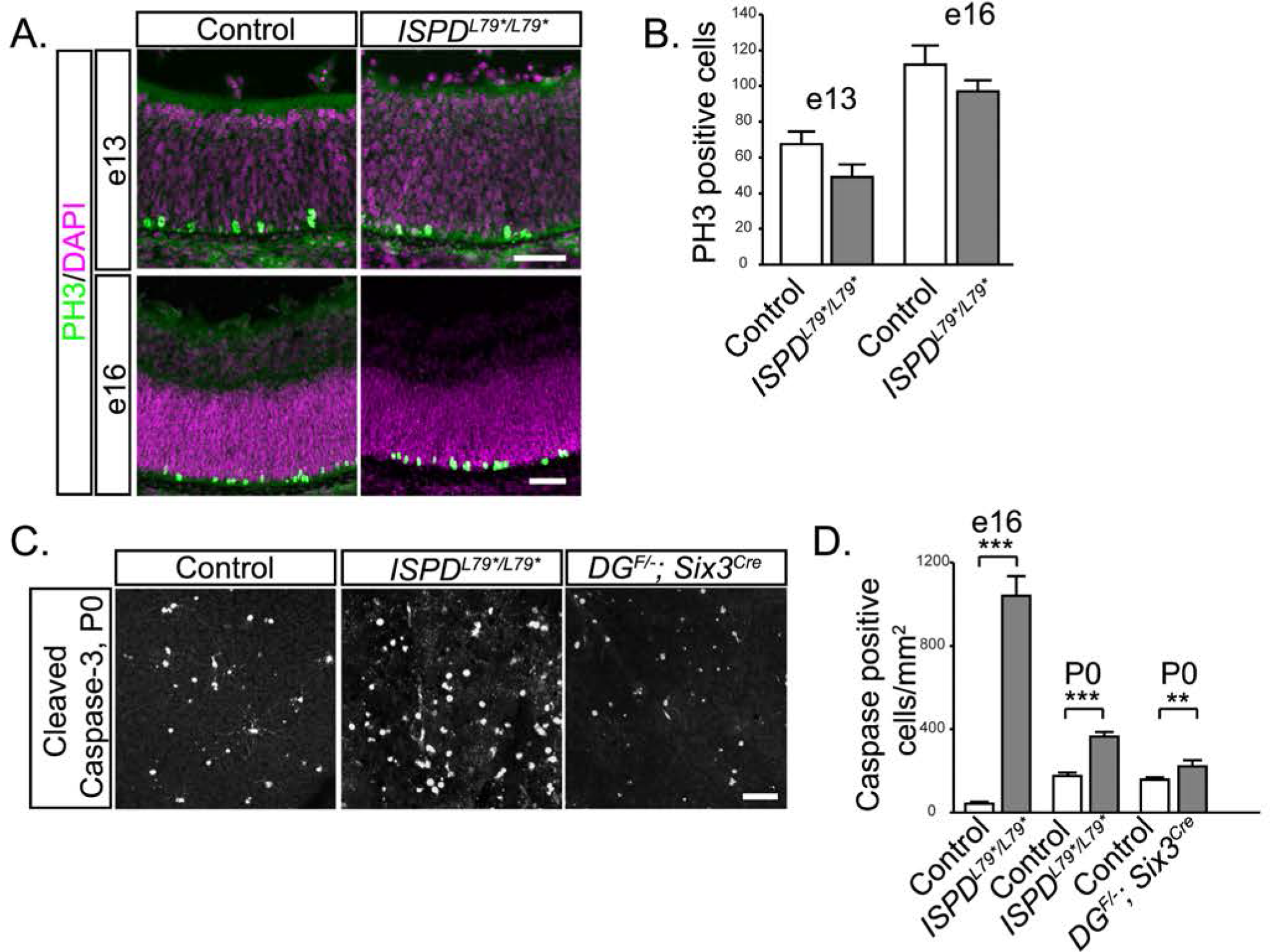
Loss of dystroglycan results in increased developmental cell death. (A) Immunohistochemistry for mitotic cells (PH3) at e13 (top) and e16 (bottom) shows normally positioned mitotic retinal progenitor cells adjacent to the RPE. (B) Quantification of mitotic cells shows no difference between control and ISPD^L79*/L79*^ – mutants (p>0.05, t test, n=4 control and 4 mutant retinas at e13, p>0.05, t test, n=4 control and 4 mutant retinas at e16). (C-D) Immunohistochemistry for cleaved caspase-3 in a flat mount preparation of P0 *DG^F/-^*; *Six3^cre^* retinas shows an increase in apoptotic cells. (p=0.0279, t test, n=18 samples from 6 control retinas, 17 samples from 6 mutant retinas). (D) Quantification of cleaved caspase-3 positive cells shows an increase in apoptotic cells at e16 (p<0.0001, t test, n=18 samples from 6 control retinas, 18 samples from 8 mutant retinas) and P0 (C, D p<0.0001, t test, n=18 samples from 6 control retinas, 15 samples from 5 mutant retinas) between control (left) and *ISPD^L79*/L79*^* (middle) retinas. Scale bar 50 μm.

### Dystroglycan functions non-cell autonomously as an extracellular scaffold in the developing retina

We next sought to provide mechanistic insight into how dystroglycan regulates retinal development *in vivo*. In the cerebral cortex, the loss of dystroglycan results in breaches of the pial basement membrane and detachment of neuroepithelial endfeet from the pial surface, depriving neurons of a migratory scaffold. In addition, the cortical basement membrane defects cause the mis-positioning of Cajal-Retzius cells, which are the source of Reelin that regulates somal translocation of neurons as they detach from the neuroepithelial scaffold (Nakagawa et al., 2015). Deletion of *dystroglycan* specifically from postmitotic cortical neurons does not result in a migration phenotype (Satz et al., 2010), supporting a model in which the cortical migration phenotypes arise due to disrupted interactions between the basement membrane and neuroepithelial scaffold. In contrast to the cerebral cortex, the basal migration of RGCs does not involve contact with the neuroepithelial scaffold. Instead, newly born RGCs migrate via somal translocation using an ILM-attached basal process that eventually becomes the nascent axon (Randlett et al., 2011; Icha et al., 2016). Dystroglycan’s expression in ILM-attached basal processes (Figure 1A) and RGCs (Montanaro et al., 1995) and the restriction of neuronal migration and axon guidance defects to the GCL raise the possibility that dystroglycan could be functioning cell-autonomously in the basal processes of newly born RGCs. To test this possibility, we generated *DG^F/-^*; *Isl1^cre^* conditional knockouts. *Islet1* is expressed in the majority of ganglion cells as they differentiate from the retinal progenitor pool (Pan et al., 2008b). Analysis of *Isl1^cre^* mice at e13 confirmed recombination occurs in the majority of newly born ganglion cells, but not in neuroepithelial progenitors (Figure 10A). Interestingly, dystroglycan protein expression in *DG^F/-^*; *Isl1^cre^* mutants was not altered compared to controls, suggesting that RGCs do not provide a significant source of dystroglycan at the ILM (Figure 10B). Examination of *DG^F/-^*; *Isl1^cre^* mutants indicated that deletion of *dystroglycan* selectively from RGCs did not affect ILM integrity (Figure 10C). Neuronal migration (Figure 10C, E and F), axon guidance (Figure 10C and D) and the stratification of dendrites in the IPL (Figures 10 E and F) all appeared normal in *DG^F/-^*; *Isl1^cre^* mutants. These results demonstrate that dystroglycan is not required within RGCs themselves during retinal development.

**Figure 10:**
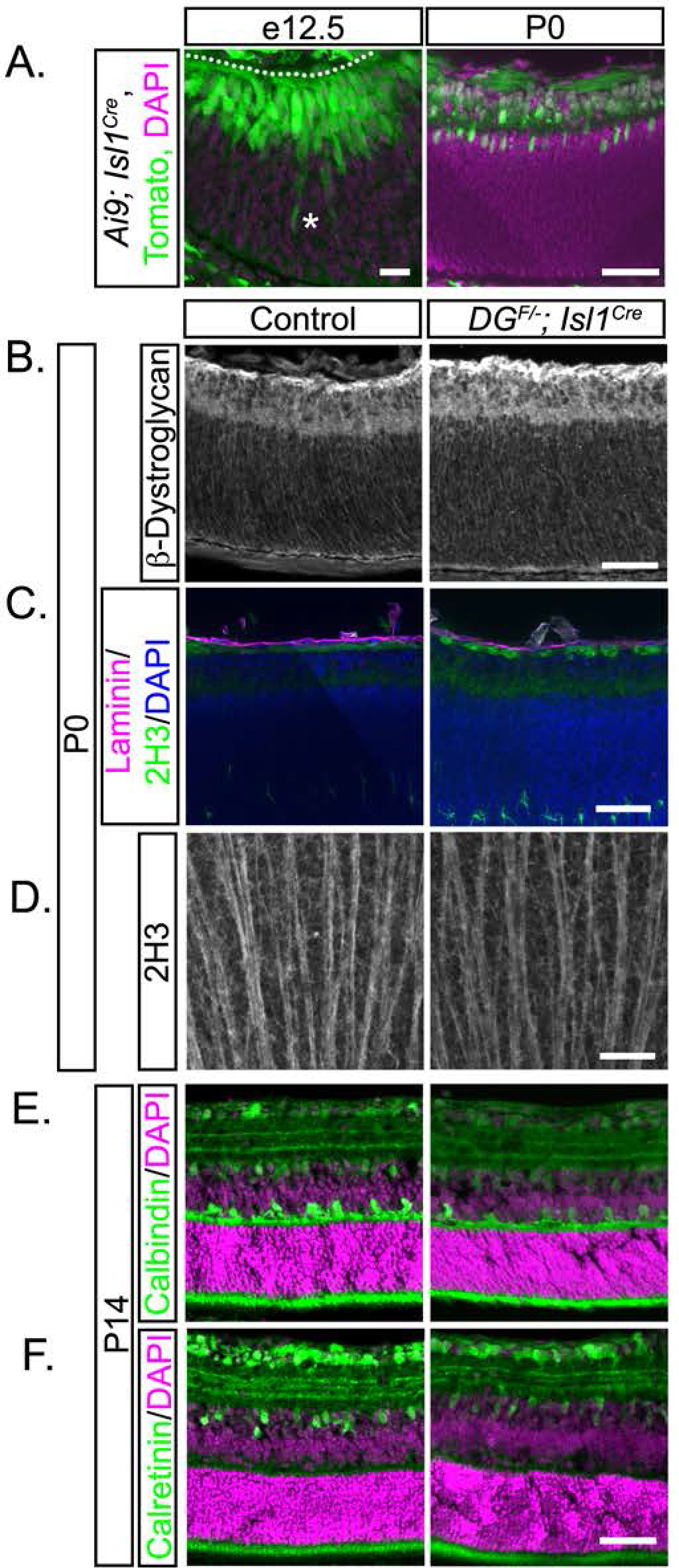
Dystroglycan is not required within RGCs for their migration and axon outgrowth. (A) Recombination pattern of *Rosa26-lox-stop-lox-TdTomato; Ai9* reporter (green) by *Isl1^cre^* by shows expression the majority of differentiated ganglion cells at e12.5 (left) and P0 (right). (B) Dystroglycan expression at the ILM is unchanged between control and *DG^F/-^*; *Isl1^cre^* retinas. (C) The ILM (laminin, purple) and axons in the OFL (2H3, green) in *DG^F/-^*; *Isl1^cre^* retinas appear similar to control. (D) Flat mount preparations show normal axon fasciculation (2H3) in *DG^F/-^*; *Isl1^cre^* retinas. (E-F) *DG^F/-^*; *Isl1^cre^* (right panel) retinas (P14) have normal cellular lamination and dendritic stratification. Dashed line indicates ILM. Asterisk notes a differentiated ganglion cell body that is still migrating toward the ILM. Scale bar 20 μm A left panel, scale bar 50 μm A right panel, B-**F**.

Dystroglycan consists of two subunits that can play distinct roles in the overall function of the protein. The extracellular α-subunit is heavily glycosylated and functions as an extracellular scaffold by binding to extracellular matrix components such as laminin. The β-subunit contains a transmembrane and intracellular domain, and can bind directly to the intracellular scaffolding protein dystrophin and other modifiers of the actin cytoskeleton, as well as initiate intracellular signaling cascades (Moore and Winder, 2010). The intracellular domain of dystroglycan is required for the localization of dystrophin to the ILM, and mice lacking the intracellular domain of dystroglycan (*DG^-/_β_cyt^*) (Satz et al., 2009) or the predominant retinal isoform of dystrophin (*Mdx^3Cv^*) (Blank et al., 1999) have abnormal scotopic electroretinograms, suggesting a defect in retinal function. While these mice do not have any disruptions in the ILM or gross malformations in the retina, whether dystroglycan signaling through dystrophin is required for neuronal migration, axon guidance or dendritic stratification has not been examined. Consistent with the original report, examination of the ILM and overall architecture of the retina is normal in *DG^-/_β_cyt^* mice (Figure 11A). In addition, we find that neuronal migration, axon guidance and stratification of dendritic lamina are unaffected in *DG^-/_β_cyt^* mice (Figure 11A-D). Therefore, intracellular signaling, including through dystrophin, is not required for these aspects of retinal development. Taken together with our results in *DG^F/-^*; *Isl1^cre^* mutants, these findings indicate that dystroglycan primarily functions within neuroepithelial cells as an extracellular scaffold to regulate the structural integrity of the ILM. The progressive degeneration of the ILM then leads to secondary defects including aberrant migration, axon guidance and dendritic stratification that primarily affect the inner retina.

**Figure 11:**
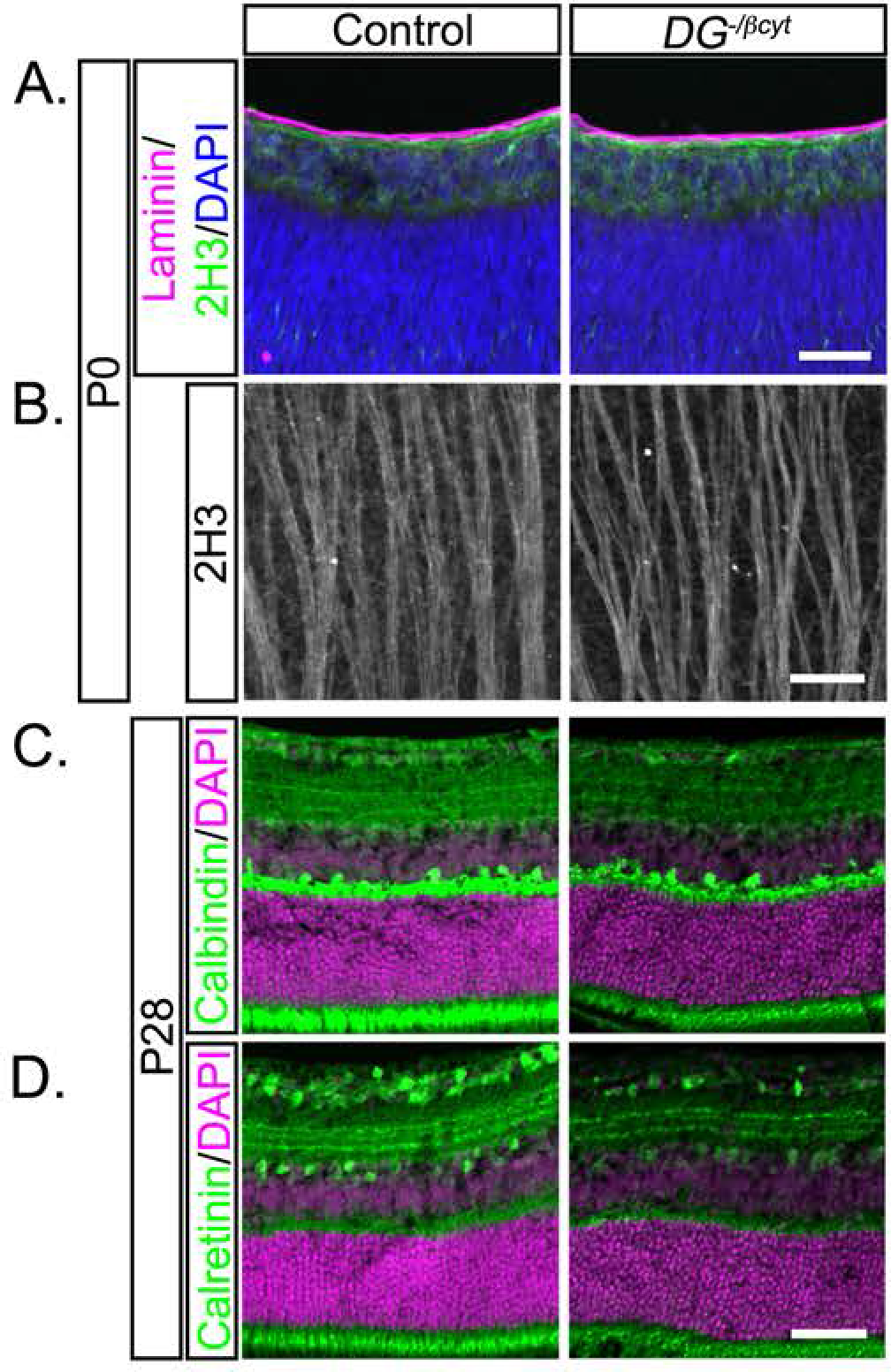
The intracellular signaling domain of dystroglycan is not required for proper retinal development. (A) ILM integrity (laminin, purple), neuronal migration (DAPI, blue) and axon outgrowth (2H3, green) all appear normal in mice lacking the intracellular domain of dystroglycan (*DG^-/_β_cyt^*) at P0. (B) Flat mount preparations show normal axon fasciculation (2H3) in *DG^-/_β_cyt^* retinas. (C-D) *DG^-/_β_cyt^* retinas (P28) have normal cellular lamination and dendritic stratification. Scale bar, 50 μm.

### Dystroglycan is dispensable for the generation of spontaneous retinal waves

One of the critical functions for laminar targeting in neural circuit development is to ensure that the axons and dendrites of appropriate cell types are in physical proximity to one another during synaptogenesis. In addition to regulating the laminar positioning of neurons in the cortex and retina, dystroglycan is required for the development of a subset of inhibitory synapses in the brain (Fruh et al., 2016). Therefore, we investigated the possibility that the loss of dystroglycan disrupts synapse formation in the retina. Previous studies have shown that ribbon synapses in the OPL are dysfunctional in the absence of dystroglycan or its ligand pikachurin (Sato et al., 2008). These synapses are normal at the resolution of light microscopy, but electron microscopy reveals that dystroglycan and pikachurin are required for the insertion of bipolar cell dendrite tips into ribbon synapse invaginations (Omori et al., 2012). Consistent with these results, we found that pre- and post-synaptic markers for ribbon synapses are present in *DG^F/-^*; *Six3^cre^* mutants (Figure 12A). In the inner retina, markers for excitatory synapses (VGLUT1, Figure 12B) and inhibitory synapses (VGAT, Figure 12C) were present in the IPL, and were also present in the mis-localized ectopic cell clusters that protrude into the vitreous (asterisks). This finding is similar to a recent study in which mice lacking the Cas family of intracellular adaptor proteins express synaptic markers localized to aberrant neuronal ectopia that protrude into the vitreous (Riccomagno et al., 2014). These results suggest that mis-laminated neurons in the retina are still able to recruit synaptic partners, despite their abnormal location.

**Figure 12:**
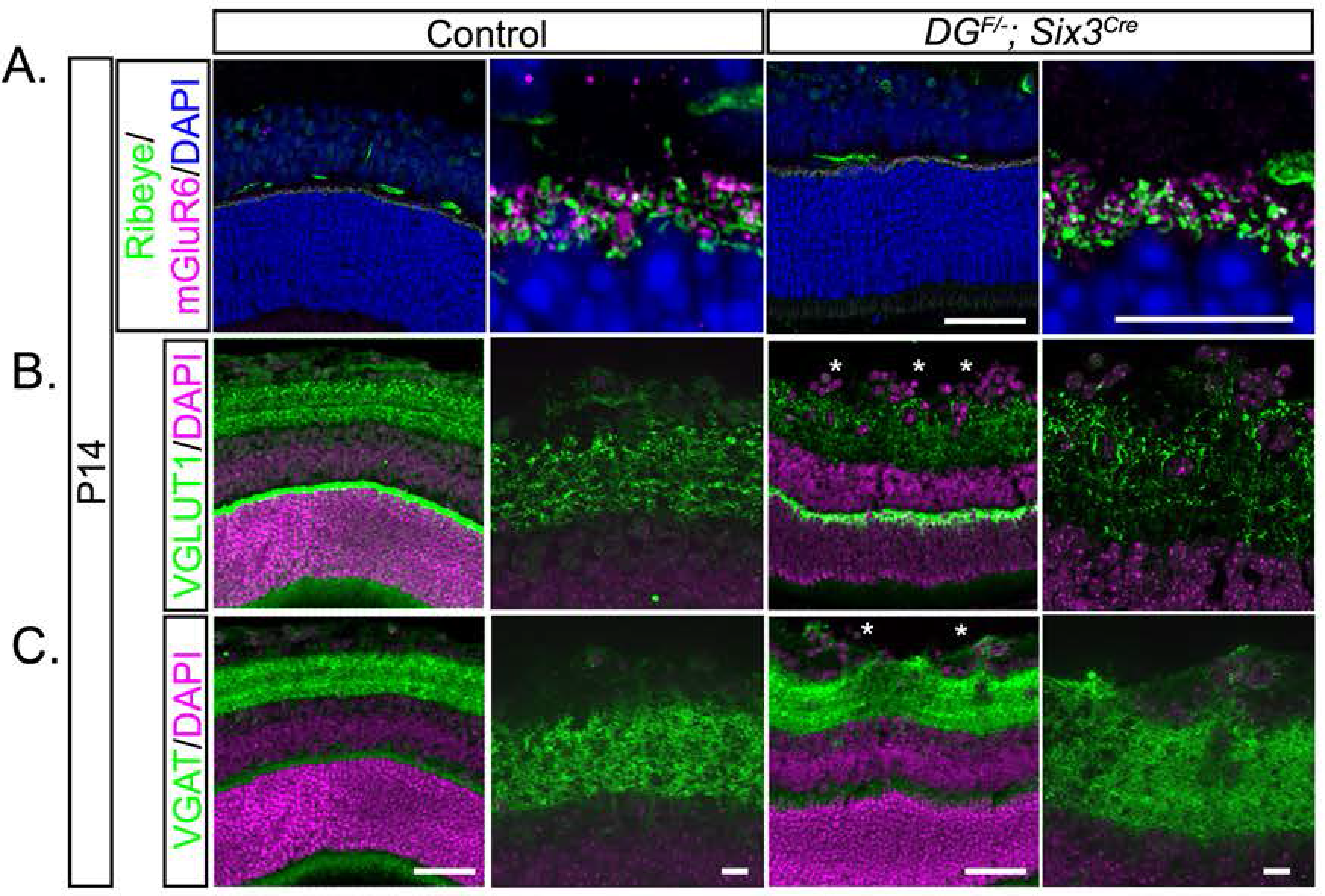
Synaptic markers are present in the retina of *dystroglycan* mutants. (A) Markers for outer retinal ribbon synapses (Ribeye, presynaptic and mGluR6, postsynaptic) appear structurally normal in the absence of dystroglycan. The density of (B) excitatory (VGLUT1) and (C) inhibitory (VGAT) presynaptic markers appear similar to control in the inner retinas of *DG^F/-^*; *Six3^cre^* mutants. Synapses are also present within ectopic clusters (asterisks). Scale bar, 50 μm wide view, scale bar, 10 μm enlarged view.

The presence of synaptic markers in ectopic neuronal clusters in the inner retina does not guarantee normal function of these neurons. Recording synaptic activity in RGCs in response to light stimuli in *DG^F/-^*; *Six3^cre^* mutants is not feasible due to the requirement for dystroglycan at photoreceptor ribbon synapses. We instead analyzed retinal waves, which are spontaneous bursts of activity that propagate across the retina prior to eye opening and are independent of light stimulation. During early postnatal development, these waves are initiated by acetylcholine (ACh) release from starburst amacrine cells and propagate along the starburst amacrine cell network prior to transmission to RGCs (Xu et al., 2016). In *DG^F/-^*; *Six3^cre^* mutants, ChAT positive starburst amacrine cells are present in normal numbers, and while they are normally localized and mosaically spaced in the INL, they are disorganized in the GCL. Therefore, we expected that these defects might affect the propagation of retinal waves through the starburst amacrine cell and RGC network. As disruptions in the ILM in *DG^F/-^*; *Six3^cre^* mutants would lead to unequal bulk loading of cell permeable calcium indicators, we utilized the genetically-encoded calcium indicator GCaMP6f crossed onto the *DG; Six3^cre^* line to visualize retinal waves.

Retinal waves in control *DG^F/+^*; *Six3^cre^*; *R26-LSL-GCaMP6f* retinas at P1-P2 were robust and had similar spatiotemporal features (area, rate of propagation, refractory period) to waves measured using cell-permeable calcium indicators (Arroyo and Feller, 2016). Consistent with previous reports, there is a broad distribution of wave area in control retinas (Figures 13A, C, Movie 1). Neighboring waves do not overlap with one another, but rather tile the retinal surface during the two-minute imaging period. To our surprise, retinal waves were present and appeared grossly normal in *DG^F/-^*; *Six3^cre^; R26-LSL-GCaMP6f* mutants (Figure 13B, Movie 2). Waves in control and mutant retinas exhibited a similar distribution in wave area (Figure 13C), and the average wave area showed no statistical difference. The rate of wave propagation showed a similar distribution between controls and mutants (Figure 13D). The average rate of wave propagation showed a small, but statistically significant, decrease (Wilcoxon Rank Sum test, p=0.0493). Thus, despite the dramatic disorganization of ChAT positive starburst amacrine cells in the GCL of *DG^F/-^*; *Six3^cre^* mutants, the generation and propagation of retinal waves persisted.

**Figure 13:**
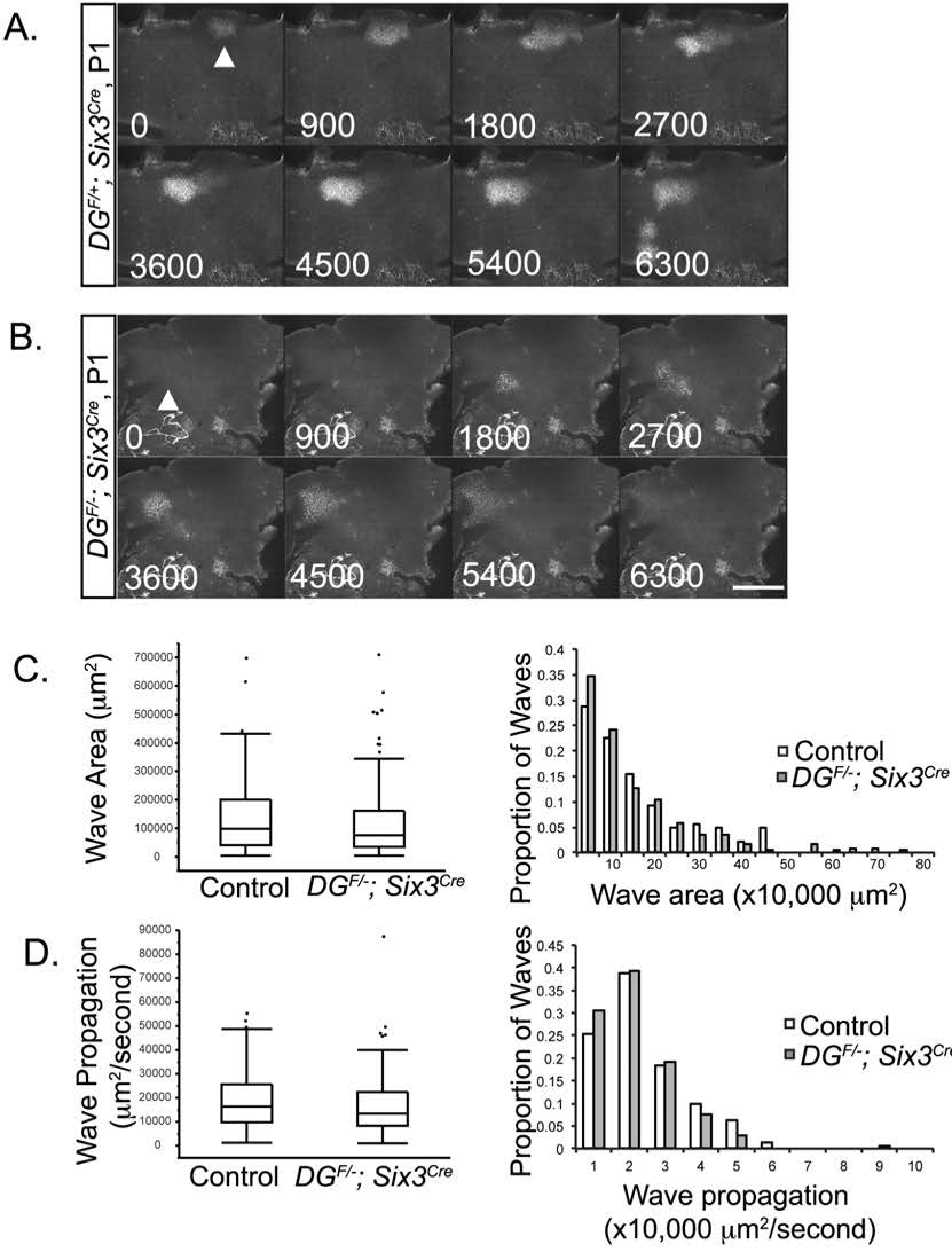
Dystroglycan is dispensable for the generation and propagation of retinal waves. (A-B) At P1, waves propagate normally across the retina in *DG^F/+^*; *Six3^cre^*; *R26-LSL-GCaMP6f* (A) and *DG^F-^*; *Six3^cre^*; *R26-LSL-GCaMP6f* mice (B). (C) Distributions of wave areas show no difference between controls and mutants (Wilcoxon rank sum test, p>0.05). (D) Wave propagation rate is slightly slower in mutant retinas (Wilcoxon rank sum test, p=0.0493). Wave parameters were calculated from 142 control waves obtained from 6 retinas from 3 control mice and 173 mutant waves obtained from 8 retinas from 5 mutant mice. Arrowheads indicate the initiation site of a retinal wave. Scale bar 500 μm. Time displayed in milliseconds.

## Discussion

While defects in retinal structure and function are observed in both human patients and mouse models of dystroglycanopathy, the mechanism of dystroglycan function in the mammalian retina and the consequence of its loss on specific cell types are poorly understood (Takeda et al., 2003; Lee et al., 2005; Satz et al., 2008; Satz et al., 2009; Chan et al., 2010; Takahashi et al., 2011). Our study establishes a critical role for dystroglycan in maintaining the integrity of the ILM, which is required for intra-retinal axon guidance and establishing laminar architecture and mosaic spacing in the inner retina. Mechanistically, we show that dystroglycan functions non-cell autonomously as an extracellular scaffold, as mice with selective deletion of *dystroglycan* from postmitotic RGCs (*DG; Isl1^cre^*) and mice lacking the intracellular domain of dystroglycan (*DG^-/_β_cyt^*) appear phenotypically normal. Despite the dramatic disruptions in cellular lamination in the GCL and dendritic stratification in the IPL in *DG; Six3^cre^* mutants, dystroglycan appears dispensable for the formation of synapses and the generation of spontaneous, light-independent activity in the retina.

### Requirement for dystroglycan at the ILM during retinal development

Using two genetic models for the complete loss of functional dystroglycan (*ISPD^L79*^* and *DG; Sox2^Cre^*), we demonstrated that after the ILM forms, it rapidly degenerates in the absence of dystroglycan. These results suggest that dystroglycan is not required for the formation of the nascent ILM, but rather plays a critical role in maintaining ILM structure as it expands to accommodate the growing retina. Recruitment of laminin is a critical early step in the assembly of the ILM, and several laminin mutants have similar disruptions in ILM integrity (Edwards et al., 2010; Pinzon-Duarte et al., 2010; Gnanaguru et al., 2013). In contrast to *Xenopus*, where depletion of *dystroglycan* results in degeneration of both the ILM in the inner retina and Bruch’s membrane in the outer retina, Bruch’s membrane is unaffected by the loss of *dystroglycan* or isoforms of *Laminin* (Lunardi et al., 2006; Pinzon-Duarte et al., 2010). Mice lacking *β-1 Integrin* in the retina exhibit a similar defect to *dystroglycan* mutants, raising the possibility that these laminin receptors may be functionally redundant (Riccomagno et al., 2014). Surprisingly however, mice in which both *dystroglycan* and *β-1 Integrin* are deleted from the retina (*DG*; *ItgB1*; *Six3^Cre^*) still formed an ILM at e13, and the subsequent degeneration of the ILM was indistinguishable from *DG*; *Six3^Cre^* mutants (unpublished observations). Therefore, sulfated glycolipids alone are likely sufficient for the recruitment of laminin during the initial formation of the ILM.

### The ILM is required for neuronal migration and axon outgrowth in the retina

In the developing cortex, the loss of functional dystroglycan leads to degeneration of the radial glia processes and ectopic clustering of reelin-secreting Cajal-Retzius neurons, which are thought to be the principal drivers of structural brain defects in dystroglycanopathies (Myshrall et al., 2012; Nakagawa et al., 2015; Booler et al., 2016). Retinal neurons do not migrate along the neuroepithelial scaffold, and there is no cue analogous to reelin to signal termination of migration, suggesting that while dystroglycan functions primarily in neuroepithelial cells in the retina, the functional implications are distinct from its role in cortical neuroepithelial cells. In the cortex of *dystroglycan* deficient mice, the organization of neurons across all lamina is affected, whereas within the retina, migration defects are restricted to amacrine cells and RGCs in the inner retina. In contrast, migration defects in dystroglycan-deficient *Xenopus* retinas are more widespread and also affect outer retinal neurons, likely reflecting the requirement for dystroglycan at both the ILM and Bruch’s membrane (Lunardi et al., 2006). What is the driving force behind the selective localization of amacrine cells and RGCs to the ectopic clusters that protrude into the vitreous in the mammalian retina? While the elimination of RGCs does not affect the lamination of other neurons, other neurons will organize themselves around mislocalized RGCs, resulting in an overall disorganization of retinal lamination (Wang et al., 2001; Kay et al., 2004; Icha et al., 2016). In *dystroglycan* mutants, RGCs that encounter the degenerating ILM and inappropriately migrate into the vitreous may then actively recruit later born neurons such as amacrine cells to inappropriate locations.

The establishment of retinal mosaics requires tangential migration that is regulated by short range interactions between immature neurites of neighboring cells (Galli-Resta et al., 1997; Reese et al., 1999; Galli-Resta et al., 2002; Huckfeldt et al., 2009). A key feature of this process is that it requires homotypic cells to be localized within the same lamina, and cells in which mosaic spacing is disrupted are no longer restricted to a two-dimensional plane (Fuerst et al., 2008; Kay et al., 2012). In *dystroglycan* mutants, the laminar organization and mosaic spacing is normal in horizontal cells and INL starburst amacrine cells, but disrupted in starburst amacrine cells in the GCL. This defect is not due to dystroglycan functioning within starburst amacrine cells, as mosaic spacing was normal in *DG*; *Isl1^Cre^* mutant retinas (data not shown). Rather, the selective defects in mosaic spacing of GCL starburst amacrine cells suggests that this is likely a consequence of disrupting the two-dimensional organization of the GCL. Alternatively, mosaic spacing of cells within the GCL may require cues present in the ILM for their tangential dispersion.

Basement membranes are highly dynamic structures that contain pro-axon growth ECM molecules such as laminins and collagens, and also regulate the distribution of secreted axon guidance cues (Halfter et al., 1987; Chai and Morris, 1999; Xiao et al., 2011; Wright et al., 2012). A number of secreted cues direct intraretinal axon guidance of RGCs. Deletion of Netrin (Deiner et al., 1997) specifically affects exit of RGC axons through the optic nerve head, and deletion of Slits (Thompson et al., 2006) or Sfrps (Marcos et al., 2015) leads to the invasion of RGC axons into the outer retina. In contrast, the randomized growth and defasciculation of axons we observed in *ISPD^L79*^* and *DG*; *Six3^Cre^* mutant retinas is more consistent with defects observed upon deletion of adhesion receptors (Bastmeyer et al., 1995; Brittis et al., 1995; Ott et al., 1998). These results suggest that dystroglycan primarily functions to organize the ILM as a substrate for axonal adhesion.

### Loss of retinal neurons in the absence of dystroglycan

While previous studies of mouse models of dystroglycanopathy have consistently noted retinal thinning, it was unclear which retinal cell types were affected. Our comprehensive analysis found that lack of dystroglycan led to reductions in photoreceptor layer thickness, horizontal cell number, and RGC number (Figure 8). Analysis of e13, e16 and P0 retinas from *ISPD^L79*^* and *DG; Six3^cre^* mutants indicated that the loss of RGCs was not due to altered proliferation of retinal progenitors, but was primarily due to increased apoptosis of cells in the GCL that preceded and extended into the normal window of developmental apoptotic cell death. This increase in apoptotic cell death did not persist in adult (P56) *DG; Six3^cre^* mutants (data not shown), suggesting it was likely restricted to the developing retina. Why are RGCs lost at such a high rate in *DG; Six3^cre^* mutants, while displaced amacrine cell number in the GCL remains normal? One possibility is that RGCs are selectively affected since they are the only cell type to project out of the retina. Indeed, we observed profound guidance defects of RGCs at the optic chiasm in *ISPD^L79*^* and *DG; Six3^cre^* mutants (unpublished observations). This result is similar to *Isl1* and *Brn3b* deficient mice, in which axon growth defects at the optic chiasm precede an increase in RGC apoptosis (Gan et al., 1999; Pan et al., 2008a). The death of RGCs whose axons fail to reach retinorecipient regions of the brain is consistent with the need for target-derived factors to support their survival, although the identity of these factors remains elusive.

While we did not observe increased caspase-3 reactivity in photoreceptors or horizontal cells at P0, it is possible that the loss of cells occurred gradually during the first two postnatal weeks. Dystroglycan is required for the proper formation of ribbon synapses between photoreceptors, horizontal cells and bipolar cells in the OPL (Sato et al., 2008; Omori et al., 2012). Therefore, the loss of appropriate synaptic contact in the absence of dystroglycan may lead to the elimination of a proportion of photoreceptors and horizontal cells.

### Persistence of retinal waves in the absence of dystroglycan

The defects in lamination and dendritic stratification of starburst amacrine cells in *DG; Six3^cre^* mutants led us to hypothesize that this would affect their ability to generate retinal waves. These waves are initiated by the spontaneous activity of starburst amacrine cells, independent of light stimuli, allowing us to circumvent the requirement for dystroglycan in proper transmission at ribbon synapses. Contrary to our expectations, retinal waves were present and propagated normally in *DG; Six3^cre^* mutants (Figure 13). The persistence of retinal waves even in the context of disrupted cellular organization supports the model that these waves are the product of volume release of ACh from starburst amacrine cells that can trigger extra-synaptic responses in cells that are not physically connected (Ford et al., 2012). Therefore, the relatively normal organization of INL starburst amacrine cells may be sufficient to overcome the disorganization of GCL starburst amacrine cells.

### Conclusion

Retinal dysplasia and optic nerve hypoplasia are frequently observed in patients with severe forms of dystroglycanopathy (Manzini et al., 2008). Using multiple mouse models, we demonstrate that dystroglycan is required for multiple aspects of retinal development. We show that dystroglycan functions within neuroepithelial cells in the retina to regulate the structural integrity of a basement membrane (the ILM), which is required for the coordination of neuronal migration, axon guidance and dendritic stratification in the inner retina. In addition, we find that there is a significant loss of photoreceptors, horizontal cells and almost 50% of RGCs due to increased apoptotic cell death. Our data suggest that the disorganization of the inner retina resulting from degeneration of the ILM is a key contributor to visual impairment in dystroglycanopathies.

## Acknowledgements

We thank Patrick Kerstein and Kylee Rosette for their technical assistance; Marla Feller and Franklin Caval-Holme for advice on visualizing and analyzing retinal waves; W. Rowland Taylor and Teresa Puthussery and members of their labs for antibodies and technical advice; David Pow for the GlyT1 antibody; Catherine Morgans for the mGluR6 antibody, Stefanie Kaech Petrie and the OHSU Advanced Light Microscopy Core (ALM) for assistance with confocal imaging; David Ginty, Alex Kolodkin, Martin Riccomagno, Randall Hand and members of the Campbell and Wright laboratories for discussion throughout the course of this study and comments on the manuscript. This work was supported by NIH Grants R01-NS091027 (K.M.W.), The Whitehall Institute (K.M.W.), The Medical Research Foundation of Oregon (K.M.W.), NSF GRFP (R.C.), LaCroute Neurobiology of Disease Fellowship (R.C.), Tartar Trust Fellowship (R.C.), NINDS P30–NS061800 (OHSU ALM), and a Paul D. Wellstone Muscular Dystrophy Cooperative Research Center grant to K.P.C. (1U54NS053672). K.P.C. is an investigator of the Howard Hughes Medical Institute.

---

Movie 1: Retinal waves in *DG^F/+^*; *Six3^cre^*; *R26-LSL-GCaMP6f* mice. 15frames/sec, total of 21 frames

Movie 2: Retinal waves in *DG^F/-^*; *Six3^cre^*; *R26-LSL-GCaMP6f* mice. 15frames/sec, total of 21 frames

